# Changes in Muscle Synergy Structure and Activation Patterns Underlie Force Field Adaptation, Retention, and Generalization

**DOI:** 10.1101/2024.12.16.628548

**Authors:** Michael Herzog, Denise J. Berger, Marta Russo, Andrea d’Avella, Thorsten Stein

## Abstract

1

Humans can adapt their motor commands in response to errors when they perform reaching movements in new dynamic conditions, a process called motor adaptation. They acquire knowledge about the new dynamics, which they can use when they are re-exposed and, limitedly, generalize to untrained reaching directions. While force field adaptation, retention, and generalization have been thoroughly investigated at a kinematic and kinetic task level, the underlying coordination at a muscular level remains unclear. Many studies propose that the central nervous system uses low-dimensional control, i.e., coordinates muscles in functional groups: so-called muscle synergies. Accordingly, we hypothesized that changes in muscle synergy structure and activation patterns represent the acquired knowledge underlying force field adaptation, retention, and generalization. To test this, 36 male humans practiced reaching to a single target in a viscous force field and were tested for retention and generalization to new directions, while we simultaneously measured muscle activity from 13 upper-body muscles. We found that muscle synergies used for unperturbed reaching cannot explain the muscle patterns when adapted. Instead, muscle synergies specific to this adapted state were necessary, alongside a novel four-phasic pattern of muscle synergy activation. Furthermore, these structural changes and patterns were also evident during retention and generalization. Our results suggest that reaching in an environment with altered dynamics requires structural changes to muscle synergies compared to unperturbed reaching, and that these changes facilitate retention and generalization. These findings provide new insights into how the central nervous system coordinates the muscles underlying motor adaptation, retention, and generalization.

**Significance Statement:** Humans can adapt their reaching movements to new dynamic conditions. They acquire knowledge about the new dynamics and can use it not only when re-exposed to these conditions but also, in part, to generalize unpracticed reaching directions. While adaptation, retention, and generalization in a force field with new dynamics have been thoroughly investigated at a kinematic and kinetic task level, coordination of the underlying muscles remains elusive. Our results show how muscle synergies - functional groupings of co-activated muscles - underlie adaptation, retention, and generalization. In particular, we observed structural changes in the muscle synergies after adaptation compared to unperturbed reaching. These changes facilitate retention and spatial generalization. Thus, muscle synergies provide new insights into human motor adaptation.

## 3 Introduction

Humans can adapt their motor commands in response to errors when their reaching movements are perturbed, a process called motor adaptation (Shadmehr and Mussa-Ivaldi, 1994). Furthermore, they can re-use the acquired knowledge when they are perturbed again and can partly generalize it to unpracticed reaching directions (Brashers-Krug et al., 1996; Gandolfo et al., 1996; Ghez et al., 1999; Shadmehr, 2004, 2017; Rezazadeh and Berniker, 2019). Many studies have thoroughly analyzed force field adaptation, retention, and generalization at the level of task-related variables; describing and modeling the mapping of end-point kinematics and kinetics (Wolpert and Kawato, 1998; Thoroughman and Shadmehr, 2000; Diedrichsen et al., 2010; Krakauer and Mazzoni, 2011). However, how the CNS coordinates motor adaptation at the level of muscle activations has not been fully investigated. The CNS may implicitly represent acquired knowledge and generate motor commands by organizing muscle synergies (Bernstein, 1967; Mussa-Ivaldi, 1999; Bizzi et al., 2008; Giszter, 2015; d’Avella, 2016). Through muscle synergies - coordinated recruitment of groups of muscles acting together as functional units - the CNS may control a small number of units rather than every muscle, thereby reducing the dimensionality of the control problem (Bernstein, 1967; Bizzi et al., 1991; Tresch et al., 1999). Furthermore, by flexibly combining and sharing synergies, a large behavioral repertoire can be generated (Mussa-Ivaldi et al., 1994; Mussa–Ivaldi and Bizzi, 2000; d’Avella et al., 2003; Ting and Macpherson, 2005; Bizzi et al., 2008).

To date, it remains unexplored how muscle synergies are related to force field adaptation, retention, and generalization; and what changes in their structure and activation patterns lead to the observed task-level kinematics and kinetics. While isometric visuomotor rotation studies (Gentner et al., 2013; De Marchis et al., 2018; Severini and Zych, 2020) showed that adaptation and spatial generalization do not require additional muscle synergies, a study by Oscari et al. (2016) study hints that force field adaptation does require additional synergies. Furthermore, studies with a few muscles’ EMGs in force field adaptation show two principal mechanisms. First, there is co-contraction of muscles acting around the elbow and shoulder joints, which decreases during adaptation but does not vanish completely (Thoroughman and Shadmehr, 1999; Milner and Franklin, 2005; Franklin and Franklin, 2021). Secondly, activity of specific muscles counteracting the force field increases, and their activation timing shifts toward the movement start (Thoroughman and Shadmehr, 1999; Huang et al., 2012; Albert and Shadmehr, 2016). Furthermore, even after a plateau in kinematic- and kinetic-dependent variables, muscle activity continues to decrease, presumably to reduce effort (Franklin et al., 2003; Huang et al., 2012). Accordingly, adaptation may be represented either by a combination of baseline reaching muscle synergies or by specific, effort-optimized muscle synergies. Either way, if muscle synergies viably represent the coordination of force field adaptation at a muscular level, we expect them to also represent retention and spatial generalization. In particular, if muscle synergies change to accommodate specific requirements for adaptation to reaching in some directions, they are a poor choice for capturing muscle patterns in unpracticed reaching directions, in line with previous narrow spatial generalization findings (Gandolfo et al., 1996; Ghez et al., 1999; Rezazadeh and Berniker, 2019).

To investigate how muscle synergies underlie force field adaptation, retention, and generalization at a muscular level, we first examined task-level variables. Accordingly, we hypothesized that (H_Task_ 1) people adapt, de-adapt, and re-adapt to the force field and that (H_Task_ 2) spatial generalization decreases with distance from the practiced movement direction. Building on this, we then analyzed muscle activation patterns and hypothesized that (H_Synergies_ 1) the muscle patterns of force field adaptation can be reconstructed by a combination of baseline reaching synergies. As this hypothesis did not hold, we hypothesized that (H_Synergies_ 2) specific muscle synergies are required. Lastly, we hypothesized that (H_Synergies_ 3) muscle synergies acquired through adaptation are only locally applicable, reflecting the narrow spatial generalization force field adaptation findings.

## 4 Material & methods

### 4.1 Participants

Thirty-six right-handed (Oldfield, 1971) male volunteers (25.9 ± 2.8 years, 1.80 ± 0.05 m height, 77.2 ± 9.5 kg mass) naïve to force field adaptation experiments gave written informed consent before participating. The Karlsruhe Institute of Technology (KIT) Ethics Committee approved the study.

### 4.2 Apparatus and task

The participants sat at a KINARM End-Point Lab with a virtual reality (VR) display (KINARM, Kingston, Canada). The chair’s height was individually adjusted so the participant sat upright, leaning his forehead against the VR frame, and had a 90° angle between the upper and forearm. In the starting position, the handle was located on the mid-sagittal plane in front of the torso. The participants performed 15cm center-out point-to-point movements in the horizontal plane with their right hand. The VR display showed the handle’s position as well as the starting and target points but obscured the view of the handle, hands, and arms. Participants were instructed to move the handle from the start to the target within 550 ± 50 ms after the handle had resided at the start point for at least 800 ms. The target color changed when reached, giving participants feedback on whether the specified movement time was met (green: within the time frame, blue: too fast, red: too slow). After the handle remained within the target for 800 ms, the manipulandum moved it back to the start for the subsequent trial. The five targets were located at -90°, -45°, 0°, 45°, and 90° (Figure 1).

**Figure 1:**
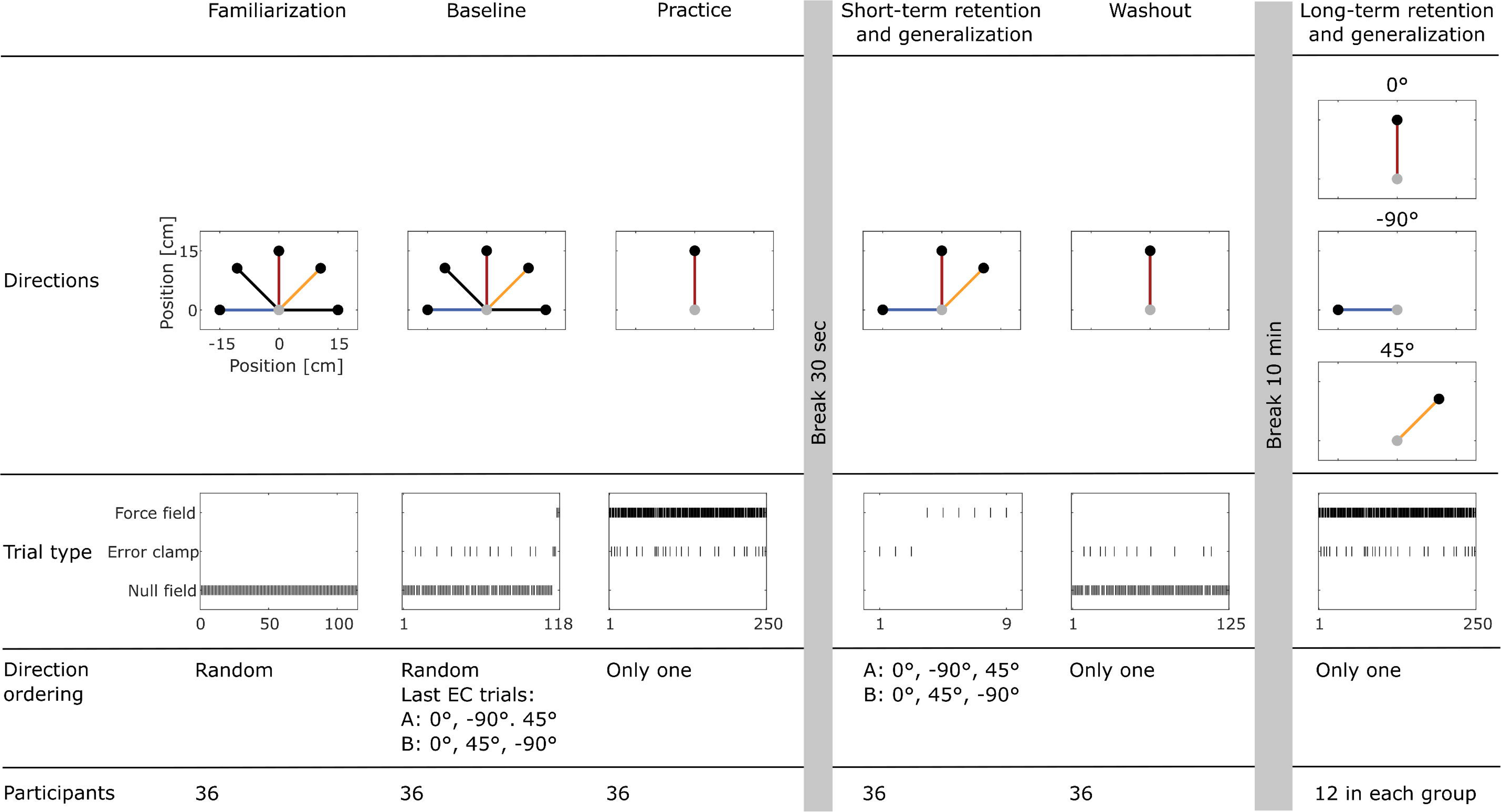
Experimental protocol. Participants performed center-out reaching movements of 15cm in length. Start points are indicated by gray circular markers and target points by black circular markers. The top row shows the movement directions, and the second row shows the sequences of trial types. After washout, the 36 participants were randomly and evenly assigned to either the long-term retention, -90° generalization, or 45° generalization group.

### 4.3 Experimental design

#### 4.3.1 Trial conditions

Three trial types were used: null field (NF), force field (FF), and error clamp (EC). In NF trials, the handle was freely movable without perturbing forces. In FF trials, a counterclockwise, velocity-dependent force field acted according to the formula:

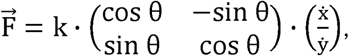

with k being the force field magnitude fixed at 20 Ns/m. The angle θ was fixed at 90°. Therefore, the force field always steered the handle’s movement orthogonally to its direction of motion. The handle’s velocity components are given by <u>^X^</u> . In EC trials, the End-Point Lab restricted the motion to a small channel connecting the start and end points (Scheidt et al., 2000; Joiner and Smith, 2008). Therefore, the manipulandum created virtual walls with a viscosity of 10 kNs/m and a stiffness of 1 kN/m.

#### 4.3.2 Groups assignment and schedule

The schedule consisted of six successive phases: familiarization, baseline, practice, short-term retention and generalization, washout, and long-term retention and generalization (Figure 1). Participants accustomed themselves to the task during the familiarization phase, which consisted only of NF trials with the directions in a random order (same order for all participants). During baseline, they reached for each target 20 times without perturbing forces (NF trials) and three times in the EC condition in a random order. The last three were FF trials in the -90°, 0, and 45° directions. During practice, they performed 250 reaches in the FF condition to the 0° target. Therein, 26 EC trials were randomly interspersed. After a 30-second break, participants were tested for short-term retention (0° target) and generalization (-90° and 45° targets). Therefore, the three targets were reached first once in the EC and then twice in the FF condition. The short-term retention and generalization started with the 0° target. Half of the participants continued with the -90° target, the other with the 45° target. This was followed by the washout, consisting of 125 trials (114 NF and 11 EC). After a 10-minute break, the long-term retention and generalization phase followed. Every participant was randomly assigned to one of three groups (-90°, 0°, or 45°; each N=12). The participants reached 250 times to one of the three targets only (target depending on group assignment) with perturbing forces. Twenty-six EC trials were interspersed at the same trial number as during the practice phase. Participants could let go of the handle during the breaks.

### 4.4 Data analysis

Kinematic (hand position and velocity) and kinetic (interaction forces) data measured at the manipulandum’s handle were recorded at 1,000 Hz with KINARM Dexterit-E software (KINARM, Kingston, ON, Canada).

Thirteen surface EMG electrodes (4,000 Hz; Noraxon USA, Scottsdale, AZ, USA) captured upper-body muscle activity of the following muscles: trapezius (descending “TrapD”, transverse “TrapT”, ascending “TrapA”), deltoid (anterior “DeltA”, middle “DeltM”, posterior “DeltP”), latissimus dorsi (“LatDorsi”), pectoralis major (“PectMaj”), serratus anterior (“SerrA”), triceps brachii (lateralis “TriLat” and medialis “TriMed”), biceps brachii (long head, “Bic”), and brachioradialis (“Bra”). Participants’ skin was prepared by shaving, abrasion, and cleansing with alcohol to ensure good electrode-skin contact. Then, Ag/AgCl electrodes were attached according to SENIAM guidelines (Hermens et al., 2000) and Perotto (2011).

#### 4.4.1 Pre-processing

Raw kinematic and kinetic data were filtered with a 4^th^-order Butterworth low-pass filter and a cut- off frequency of 6 Hz (kinematic) and 10 Hz (kinetic) following previous studies (Stockinger et al., 2015; Herzog et al., 2022). Movement start was defined as the instant the participant left the start point, and movement end when he reached the target point for the first time. Raw EMG data were bandpass filtered with a 20-450 Hz, 4^th^-order zero-lag Butterworth filter (Albert and Shadmehr, 2016). ECG artifacts apparent in the recordings of the trunk muscles were removed with a template-matching procedure (Peri et al., 2021). Subsequently, 50 Hz noise was removed with a 2^nd^-order zero-lag Butterworth notch filter (50 Hz and harmonics up to 500 Hz; Ahmad et al. 2013; Anwar et al. 2011). The filtered EMG data were full-wave rectified and envelopes were calculated using a 4^th^- order Butterworth low-pass filter with a cut-off frequency of 10 Hz. EMG data were segmented from 200 ms before the participant left the start point until 200 ms after they reached the target for the first time, e.g., including potential overshooting corrections. The segmented data were time-normalized to 101 time points. Then, the data were amplitude-normalized per muscle and participant to the maximum activity across all trials. Finally, the tonic EMG component was removed from all trials (d’Avella et al., 2006). To do so, a linear ramp was modeled using the average EMG envelope of each muscle in the 200 ms before the movement start and the 200 ms after target reach. Each calculation of averages included all baseline trials to the same direction. This direction-, muscle-, and participant-specific estimation of the tonic component was then subtracted from all respective trials. The remaining phasic parts of the EMG could contain negative values, which were clipped to zero. After careful observation, TrapD was excluded as it contributed only noise. All processing and analysis steps were performed in Matlab (R2023b, Natick, MA, USA).

#### 4.4.2 Kinematic and kinetic dependent variables

Following previous studies, adaptation was assessed with a kinematic and a kinetic measure (Sing et al., 2009; Heald et al., 2018). The kinematic variable PD_max_ quantifies the maximum perpendicular distance between a trial’s trajectory and a virtual straight line connecting the start and target. While it quantifies the net motor output with all control processes involved (Stockinger et al., 2015), the kinetic force field compensation factor (FFCF) quantifies the participant’s force field prediction (Scheidt et al., 2000; Joiner and Smith, 2008). FFCF was calculated by linear regression according to the formula:

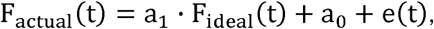

where the error e(t) was minimized (least-squares). The regression coefficient a_0_ was the axis intercept, and a_1_ was the slope. The slope serves as FFCF. F_acutal_ was the force with which the participant pressed the handle against the virtual wall. F_ideal_ was the product of the force field matrix and the trial’s velocity profile, resulting in the force profile with which the participant would have produced a straight trajectory in the force field. If F_ideal_ and F_actual_ are identical, the FFCF is 1; if unrelated, the FFCF is 0.

To consider only changes based on adaptation and generalization, all PD_max_ and FFCF values were participant- and target-specific baseline-subtracted (Wagner and Smith, 2008).

#### 4.4.3 Extraction and fitting of muscle synergies

We set up three hypotheses to investigate how force field adaptation, retention, and spatial generalization are represented in a modular structure, using the following “extract-and-fit” approach (Figure 2). To test H_Synergies_ 1, that the muscle patterns of force field adaptation can be reconstructed by a combination of baseline reaching synergies, we extracted muscle synergies from the baseline trials (see 4.4.3.1) and tested their ability to explain the muscle patterns of the adapted state by fitting them on adapted state trials (see 4.4.3.2). To test H_Synergies_ 2, that specific muscle synergies are required, we extracted shared-and-specific muscle synergies of the baseline and adapted state (see 4.4.3.3). A shared-and-specific muscle synergy extraction approach stems from the observation that synergy combinations span specific subspaces in muscle activation space, and that different sets of synergies may span the same subspace, as they can result from a rotation within the subspace. Thus, the approach aims to best identify the intersecting (i.e. shared) and the disjunct (i.e. specific) parts of the two subspaces spanned by two sets of muscle synergies (Cheung et al., 2005; d’Avella and Bizzi, 2005).

**Figure 2:**
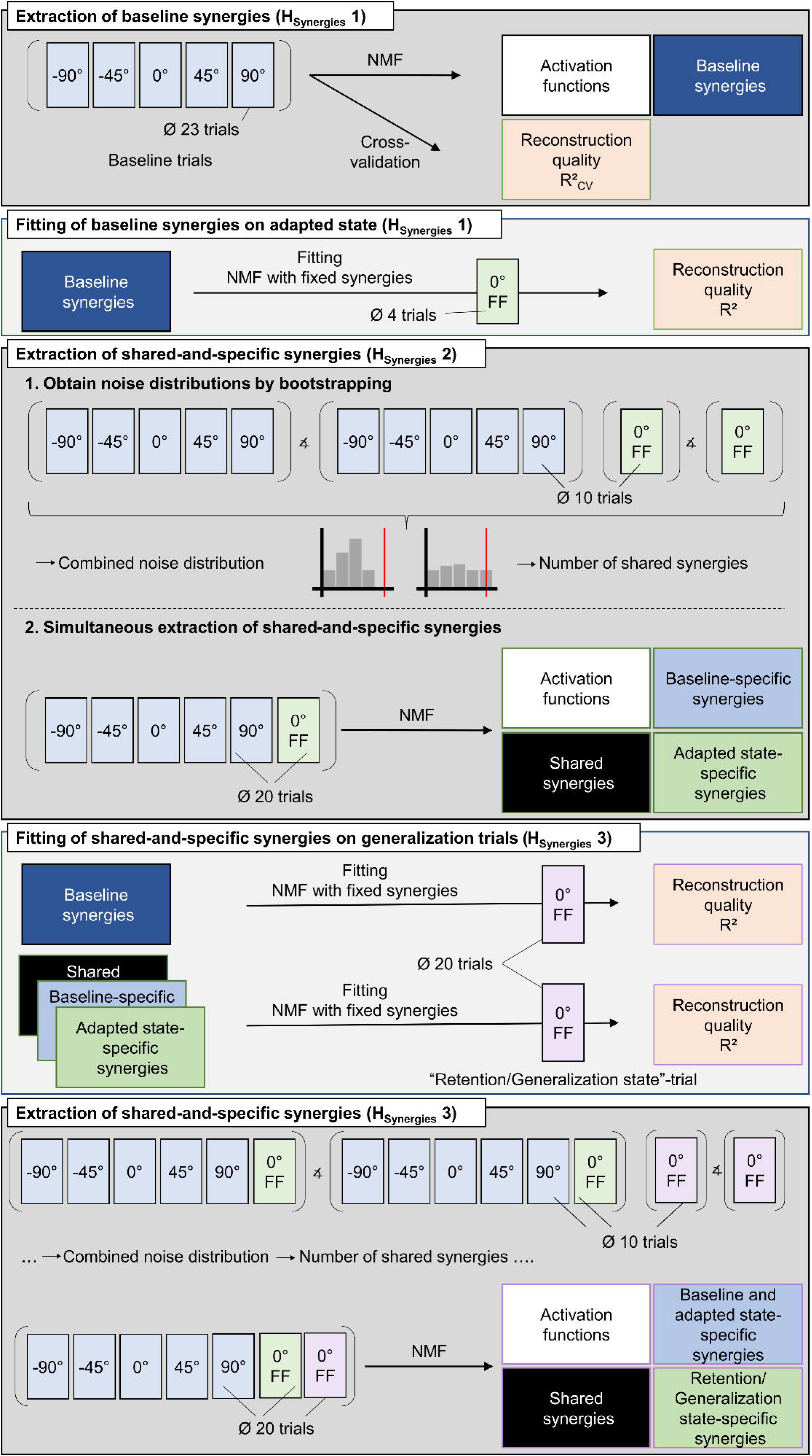
Summary of muscle synergy analyses. We followed an iterative “extract-and-fit” approach. First, baseline synergies were extracted and fitted to the adapted state. Next, shared-and-specific synergies of the baseline and adapted state were extracted and fitted to the first generalization trials. Finally, shared-and-specific synergies were extracted from the baseline and adapted state together and the first generalization trials. The ∡ sign illustrates the calculated principal angle.

Alternatives to the shared-and-specific extraction have the following limitations. First, extracting from the pooled EMG would yield a muscle synergy representation underlying the two datasets, assuming that dimensions are shared, though this assumption is what is to be tested. Second, separate extractions with a subsequent similarity analysis may be misleading as the similarity value is not unambiguous but depends on the single sets muscle synergy extraction (Cheung et al., 2009). Finally, to test H_Synergies_ 3, that muscle synergies acquired through adaptation are only locally applicable, reflecting the narrow spatial generalization force field adaptation findings, we fitted the shared-and-specific muscle synergies to the first 20 retention/generalization trials (4.4.3.4). Subsequently, we extracted shared-and-specific synergies from the baseline and adapted state together and the first retention/generalization trials (see 4.4.3.5), with the aim of identifying a muscle synergy representation of what facilitates retention and generalization.

##### 4.4.4.1 Extraction of baseline muscle synergies

For every participant, a matrix 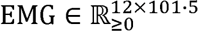 was composed, with the EMG data of the 12 muscles in rows and the five averaged trials with 101 time points each in columns. Each averaged trial consists of the averaged EMG data of the 23 baseline trials to the same target. We extracted spatial muscle synergies with non-negative matrix factorization (NMF; Lee and Seung 1999, 2001; Russo et al. 2024). NMF reduces the dimensionality of the EMG dataset by approximating it with N trial-invariant spatial muscle synergies 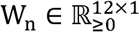, vectors specifying relative muscle activation levels, as well as N synergy activation profiles 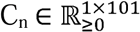:

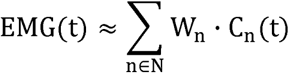

The decomposition was repeated 50 times with random initial conditions to avoid convergence to local minima and was limited to 3,000 iterations (Bach et al., 2021; Carey et al., 2021). The reconstruction quality was assessed using the multivariate R² = 1 − SSE/SST. SSE was calculated as the sum of the squared residuals and SST as the sum of the squared residuals from the mean vector (d’Avella et al., 2006). The number of extracted synergies N was chosen at the R² knee point, after which the R² curve remained approximately straight. The R² knee point was calculated using a series of linear regressions fitted to the R² versus N curve (Matlab *polyfit*), beginning with a regression across the interval [1, 12]. We then iteratively excluded the smallest value from the regression interval. We identified the optimal number of synergies N as the first N with a regression line from N to 12 with a mean square error smaller than 10^-4^ (d’Avella et al., 2006).

##### 4.4.4.2 Quality of reconstruction of adapted state data with baseline muscle synergies

To investigate if baseline reaching muscle synergies can reconstruct the muscle patterns of force field adaptation, they were tested for their ability to explain the muscle patterns (R²) in the adapted state. In particular, NMF was applied to the averaged EMG data from the last four force field trials of the adaptation phase, keeping the baseline synergies 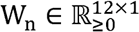 fixed and updating only the activation profiles 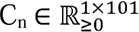. Then, we compared the R² values of the reconstruction with a cross-validated R²_CV_ of the baseline extraction. Therefore, we repeated the following process 100 times. Four randomly selected baseline trials to the same direction were averaged and constituted the test part of the cross-validation. The remaining 19 trials were averaged and horizontally concatenated with the remaining four average baseline trials, constituting the training part of the cross-validation (see 4.4.3.1). Muscle synergies were extracted from the training part and fitted to the test part. Cross-validation was used to prevent a misleading overestimation of the synergies’ reconstruction ability on the baseline phase (Stone, 1974).

##### 4.4.4.3 Extraction of shared-and-specific muscle synergies of baseline and the adapted states

To investigate if specific muscle synergies are required for force field adaptation, we extracted three sets of synergies from the combined baseline and adapted state data: one set shared between baseline and adapted state data, a second set specific to the baseline data, and a third set specific to the adapted state data (Cheung et al., 2005; d’Avella and Bizzi, 2005). We determined the number of shared synergies as the dimension of the shared subspace spanned by the synergies extracted separately from baseline and adapted states. To estimate such a dimension, we used a bootstrapping procedure that differentiated noise and structural differences (Sylos-Labini et al., 2020; Brambilla et al., 2023). This procedure assumes that differences in the synergies extracted from different subsets of trials in the same conditions are due to noise rather than structural differences.

The procedure consisted of four steps. First, muscle synergies were extracted from the baseline and adapted state separately following the description in section 4.4.3.1. Thereby, the EMG data matrix of the baseline state consisted of the 12 muscles in rows and the five averaged trials in columns. Each trial was the average of the 20 NF baseline trials to the same direction. The adapted state matrix consisted of 12 muscles in rows and one averaged trial over the last 20 FF. The principal angles between the subspaces spanned by the two sets of muscle synergies were calculated (θ_Baseline vs. adapted state;_ Golub and Van Loan 1989). Secondly, both baseline and adapted state datasets were split into two disjunct, equal-sized sub-datasets. The two baseline sub-datasets comprised five averaged trials, each from 10 randomly selected trials to the same direction. The two adapted state sub-datasets constituted one averaged trial, calculated on 10 randomly selected trials of the last 20 FF trials during adaptation. Muscle synergies were extracted from each subset, and the principal angles between the subspaces spanned by the muscle synergies of the same phase were calculated. The second step was repeated 500 times, each time with a random drawing with replacement of trials included in the subsets. This resulted in two principal angle distributions, one for the baseline and one for the adapted state. These distributions quantify the noise inherent in the baseline and the adapted state data. Thirdly, the principal angles from each subset were used to obtain a new distribution of principal angles between baseline and adapted state synergies due only to noise. To do so, two new subspaces representing the “noise subspaces” for the baseline and adapted state were calculated starting from a common base and rotating each subspace according to the respective principal angle distribution. Then, the principal angles between the two newly constructed “noise subspaces” of the baseline and adapted state were calculated. The third step was repeated 500 times, providing a distribution of the principal angles from the “noise subspaces”. Fourthly, the latter distribution’s 95^th^ percentile (θ_95_) was calculated and compared to the principal angles between the original baseline and original adapted state synergies (step one, θ_Baseline vs. adapted state_). The number of shared synergies was defined as the number of principal angles between the baseline and the adapted state, which were smaller than the corresponding θ_95_. This is based on the assumption that structural differences between muscle synergies are represented by much larger principal angles than those due to noise.

After these steps, which resulted in the number of muscle synergies shared between the baseline and adapted state, shared-and-specific muscle synergies were simultaneously extracted with an iterative process. The horizontal concatenation of the baseline and adapted state datasets, as described in the first step, constituted the input matrix for the NMF algorithm. Starting with the number of shared synergies, muscle synergies were extracted. The reconstruction qualities R²_baseline_ and R²_adapted state_ were calculated and compared to the original R² values obtained during the first step of the shared extraction. If the original R² values were not reached, a new NMF decomposition was performed with the number of shared synergies plus one or two, depending on the R² comparisons. Additionally, the matrix C with the synergy activation profiles was provided. C was initialized with random values in the cells for the shared synergies, baseline- and adapted state-specific synergies to extract. The remaining cells were filled with zeros. Therefore, using the properties of the NMF multiplicative update rule, activation profiles specific to one phase were not considered for the other phase.

##### 4.4.4.4 Fitting shared-and-specific muscle synergies of baseline and the adapted states to the first trials of the long-term retention and generalization phase

To test H_Synergies_ 3, that muscle synergies acquired through adaptation are only locally applicable, reflecting the narrow spatial generalization force field adaptation findings, we fitted (1) the baseline synergies and (2) the shared-and-specific muscle synergies extracted from baseline and the adapted state (4.4.3.3) to the average of the first 20 FF trials of the long-term retention and generalization phase and tested their ability to reconstruct the respective muscle pattern (R²). If the hypothesis holds, then the reconstruction using the shared-and-specific synergies compared to using baseline synergies would be best for the 0° group (retention), higher than zero for the 45° group (limited generalization) and zero for the -90° group (no generalization).

##### 4.4.4.5 Extraction of shared-and-specific muscle synergies of the baseline and the adapted state together and the start of the long-term retention and generalization phase

With a similar approach to the four steps described in 4.4.3.3, we extracted shared-and-specific synergies from the baseline and adapted state together and the first 20 FF retention/generalization trials. The aim was to identify a muscle synergy representation of what facilitates retention and generalization. For the EMG data matrix of the baseline and adapted state, the same baseline matrix as described in 4.4.3.3 was concatenated horizontally with one averaged trial over the last 20 FF trials during adaptation. For the EMG data matrix of the retention/generalization trials, the first 20 FF trials during this phase were averaged. We then calculated the noise distributions in a similar way to 4.4.3.3, with subsets of 10 averaged trials. We ensured that the muscle synergies extracted from the baseline and adapted state kept their structure as extracted in 3.4.3.3, i.e., their number of shared and specific synergies. Therefore, we initialized the C matrix with random numbers in cells according to the shared and specific synergies and zeros otherwise. Hence, in these calculations, the synergy vectors’ values and the activation functions’ values could vary, but the general structure of the synergy extraction remained. The combined shared-and-specific extraction in the last step also used a specific initialization of the C matrix. C was initialized with random values in the cells for the shared and specific synergies between the baseline and adapted state together and the first retention/generalization trials and zeros otherwise.

##### 4.4.4.6 Clustering of similar muscle synergies

To compare synergies extracted from different participants, we grouped them using cluster analysis. Each synergy was normalized to the maximum of its elements. First, hierarchical clustering (Matlab *pdist* (Minkowski distance; p = 3), *linkage* (Ward option), and *cluster*; Allen et al. 2019) was used to determine the optimal number of clusters. The number was chosen by assessing (1) the scree plot, which plots the within-cluster sum of squares of the linkage distance against the number of clusters, and (2) the silhouette method (Matlab *silhouette*), which quantifies the similarity of a given synergy to the other synergies in its cluster with respect to synergies from other clusters. The silhouette value s_syn_ for the synergy syn is calculated as follows:

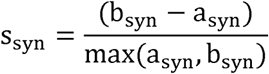

Here, a_syn_ is the average distance (cosine similarity) from syn to the other synergies of the same cluster, and b_syn_ is the minimum average distance (cosine similarity) from syn to the synergies in a different cluster, minimized over the clusters. Silhouette values range from -1 to 1, with higher values indicating a better similarity. The number of clusters was chosen at the knee point of the scree plot and increased when a cluster contained low negative values (< -0.3).

Secondly, the synergies were clustered with k-means++ based on their cosine similarity. The centroids are the synergies with the least distance (1 - cosine) to all synergies within one cluster (Sylos-Labini et al., 2020).

#### 4.4.4 Cosine tuning of the baseline synergy clusters

Synergy tuning curves were calculated for each baseline synergy cluster by a cosine fit using the integral under the mean activation of each synergy cluster and the target position (-90°, -45°, 0°, 45°, 90°; d’Avella et al. 2006). Therefore, the integrals were fitted with a linear regression according to the following formula (Matlab *regress*):

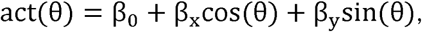

where act(Θ) is the integral under the mean activation of each synergy cluster toward the target in the direction of Θ. Synergies can be used to decelerate a movement going in the direction opposite to its “acting direction”. Therefore, synergy tuning curves were calculated for the first 25% of the movement duration, ensuring tuning only to the “acting direction”. The quality of the fits was assessed using R², and significance of the cosine tuning was assumed when the p-value of the regression between the data and the optimal cosine tuning was smaller than 0.05.

### 4.5 Statistical Analysis

#### 4.5.1 Kinematic and kinetic dependent variables

Differences in the kinematic and kinetic dependent variables, PD_max_ and FFCF, were tested as follows. Normality was assessed with Shapiro-Wilk tests and the homogeneity of variances with Levene’s tests.

Whether the participants adapted and washed out (H_Task_ 1) was tested separately with dependent t-tests comparing the means of the first and last two FF trials and the first and last EC trials, respectively.

The short-term retention and generalization (H_Task_ 2) were assessed as follows: first, a one-way repeated-measurements ANOVA with subsequent *post-hoc* t-tests identified differences between the three groups (-90°, 0°, 45°) on the PD_max_-mean of the two FF trials. Second, the FFCF values were tested for differences against zero (zero = no retention/generalization) with one-sample t-tests separately for each direction. Third, FFCF differences between the groups were tested with a one- way repeated-measurements ANOVA with subsequent *post-hoc* t-tests.

To determine if people (re-)adapt in the long-term retention and generalization phase, the three groups were tested separately with dependent t-tests on (1) the first and last two FF trials (PD_max_) and (2) the first and last EC trials (FFCF). Furthermore, the first FFCF values were tested for differences against zero (zero = no retention/generalization) with one-sample t-tests to assess retention and generalization.

To investigate H_Task_ 2, that generalization decreases with distance from the practiced movement direction, a linear mixed model (LMM) was used (Matlab *fitlme*). LMM allows the consideration of repeated measures (level 1) of a single participant (level 2), instead of the required aggregation of data as in t-tests or ANOVAs, which suits an adaptation experiment featuring inter-trial changes. The group assignment was included as a dummy variable using reference coding (either DIR90 or DIR45 set to 1, or both to 0 for the DIR0 group). The first LMM was calculated with DIR0 as the reference group and the second LMM with DIR90 as the reference group, allowing investigation of all pairwise comparisons. LMM further allows the inclusion of both FF and EC trials into the model and also the interaction of trial type with trial duration; this improved the model fit based on the change in the -2 log-likelihood and the Akaike’s information criterion (Matlab’s *linearmixedmodel.compare* function; Russell and Haworth, 2014; Ippersiel et al., 2021). The residual plots were inspected to assess linearity and homoscedasticity as prerequisites for LMM, and no gross violations were found (Hox et al., 2017).

The LMM regression formula was: 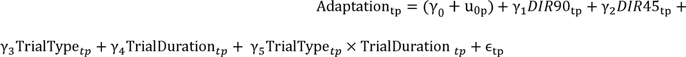 Adaptation_tp_ represents the PD_max_ or FFCF value of the t^th^ trial for the p^th^ participant. The variable u_Op_ is a participant-specific random component and the γs are fixed effect parameters for the group assignment (dummy coding), trial type, trial duration, and the interaction of trial type and duration. Accordingly, the following formula was used for the function specification of the *fitlme* function: Adaptation ∼ DIR90 + DIR45 + TrialType * TrialDuration + (1 | Participant). The LMM was implemented using the maximum likelihood method. Twenty trials were included.

#### 4.5.2 Muscle synergy analyses

A dependent t-test was used to determine differences in the reconstruction quality of the baseline muscle synergies (R²_CV_) and their fit to the adapted state (H_Synergies_ 1; section 4.4.3.2). The Kruskal-Wallis and Wilcoxon tests were used to test for improvements in the reconstruction quality between fitting the baseline synergies and fitting the shared-specific synergies on the retention/generalization trials, as the distribution was not normal (H_Synergies_ 3).

#### 4.5.3 General procedure

For all statistics, the significance level (two-tailed) was set *a priori* at 0.05. It was adjusted for multiple comparisons *post-hoc* with the Holm-Bonferroni correction (Holm, 1979), and to 0.025 for the LMM statistics as the test was carried out with two reference groups. The effect sizes were determined with η_p_^2^ and Cohen’s |d|; and classified as small (η_p_^2^ ≥ 0.01, |d| ≥ 0.2), medium (η_p_^2^ ≥ 0.06, |d| ≥ 0.5), and large (η_p_^2^ ≥ 0.14, |d| ≥ 0.8; Cohen, 1988).

## 5 Results

We were interested in how muscle synergies reflect force field adaptation, retention, and generalization at a muscular level. Therefore, we first examined whether all participants adapted, washed out, and showed retention/generalization (H_Task_ 1 and H_Task_ 2, Figure 3) based on the kinematic and kinetic dependent variables (PD_max_ and FFCF). Then, we examined the underlying muscle synergies of adaptation, retention, and generalization (H_Synergies_ 1-3).

**Figure 3:**
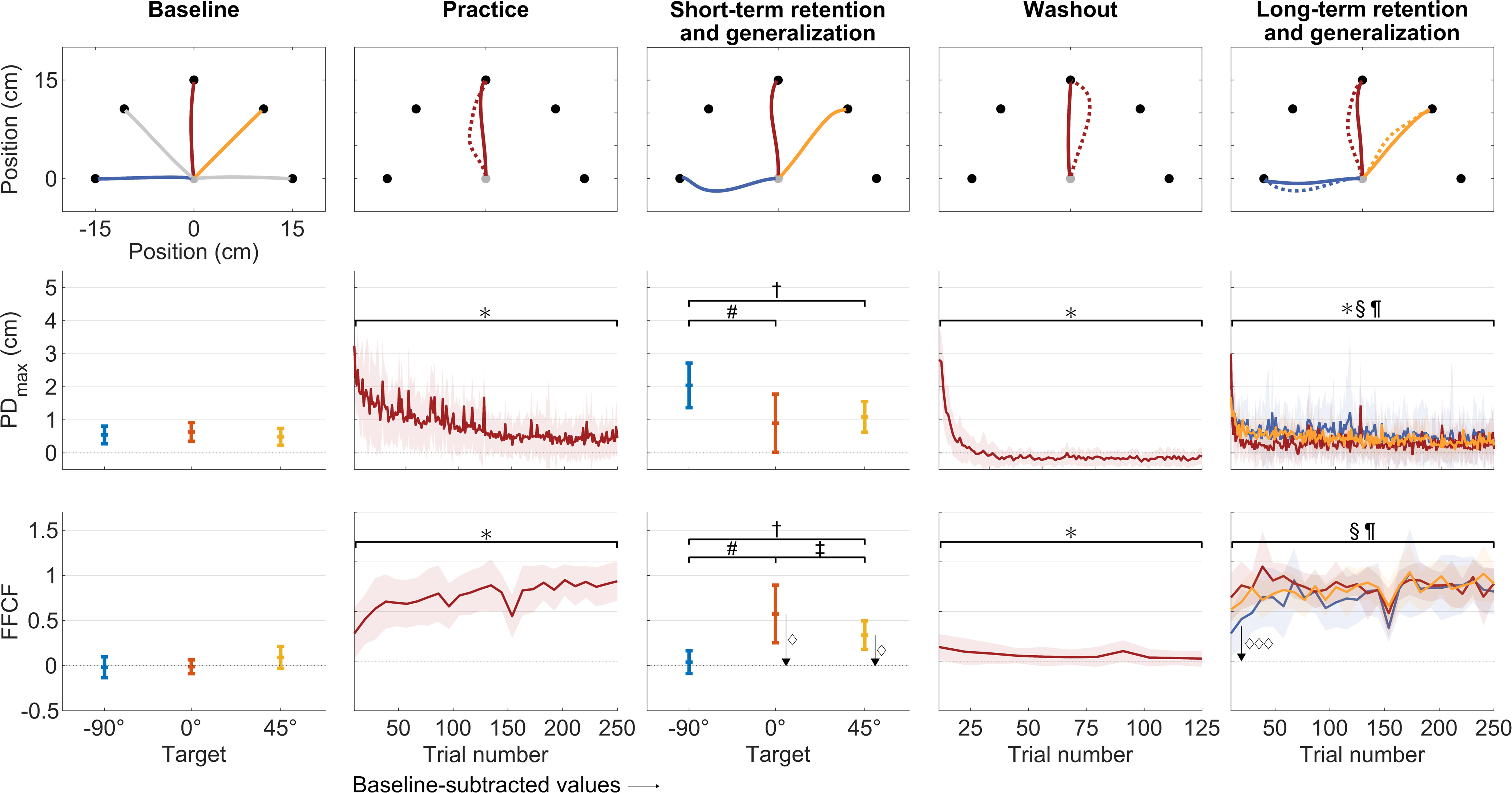
Kinematic and kinetic results. In columns: the different phases are arranged chronologically from left to right. Top row: Mean trajectories. Solid lines in baseline and short-term retention and generalization are mean values across all NF and FF trials and all participants. Dashed lines in practice, washout, and retention/generalization show mean values across all participants’ first two trials in the respective phases. Solid lines, likewise, for the last two trials. Middle and bottom rows: PD_max_ and FFCF mean and standard deviation values. The signs ∗, §, and ¶ indicate statistically significant differences over time for the 0°, -90°, and 45° targets, respectively. The # indicates a statistically significant difference between performance for the -90° and 0° target, the † between the -90° and the 45° and the ‡ between the 0° and 45° target. The l1 indicates a statistically significant different value from zero.

### 5.1 Participants adapted to the force field and washed out successfully

When first exposed to the force field, participants’ trajectories became typically curved and then, with practice, became almost straight again ([PD_max_: t(35) = 14.03, p < 0.001, d = 2.86; FFCF: t(35) = -9.15, p < 0.001, d = -2.28], Figure 3). Therefore, we can state that participants adapted to the force field. At the beginning of the washout, participants’ trajectories became curved again, mirroring the initial trajectories of the adaptation phase, and then became almost straight again [PD_max_: t(35) = 22.60, p < 0.001, d = 5.15; FFCF: t(35) = -9.15, p < 0.001, d = -2.30]. Hence, participants washed out. We therefore accept H_Task_ 1, that the participants adapt to the force field and de-adapt after its removal.

### 5.2 Participants generalized better to the 45° than to the -90° target at the short-term retention and generalization test

After the adaptation period, all participants were tested for retention and generalization to the -90° and 45° targets. The ANOVA [F(2, 58.01) = 216.45, p < 0.001, η ^2^ = 0.86] and *post-hoc* t-tests on the PD_max_ values revealed no difference between the 0° and the 45° target [t(35) = -1.59, p = 0.120, d = -0.28] but indicated that there were differences between 0° and -90° [t(35) = 8.79, p < 0.001, d = 1.58] as well as between the -90° and 45° targets [t(35) = 8.22, p < 0.001, d = 1.56]. Hence, the generalization to the 45° was not worse than the 0° retention, and better than the generalization to the -90° target.

The FFCF values were significantly different from zero for the 0° [t(35) = 10.77, p < 0.001, d = 1.76] and 45° target [t(35) = 12.91, p < 0.001, d = 2.11] but not the -90° target [t(35) = 1.84, p = 0.075, d = 0.30], indicating retention (0° target) and generalization to the 45° target only. The ANOVA [F(2, 50.32) = 53.37, p < 0.001, η_p_^2^ = 0.60] and *post-hoc* t-tests on the FFCF values showed the best performance for retention, and better performance (i.e., better generalization) for the 45° than for the -90° target [-90° vs. 0°: t(35) = -8.86, p < 0.001, d = -2.15; -90° vs. 45°: t(35) = -9.43, p < 0.001, d = -2.06; 0° vs. 45°: t(35) = 4.01, p < 0.001, d = 0.91].

In summary, in the short term, participants showed retention, and also generalized better to the 45° than the -90° target.

### 5.3 Participants re-adapted to the practiced direction and showed better generalization to the 45° than to the -90° target at the long-term retention and generalization test

After the successful washout and a 10-minute break, the participants were randomly assigned to one of the three groups (-90°, 0°, 45°) and tested if they adapted (again) and presented differences across the directions.

All groups showed retention or generalization on the first trial as assessed with t-tests vs. 0 on the FFCF values [-90°: t(11) = 5.47, p < 0.001, d = 1.47; 0°: t(11) = 13.61, p < 0.001, d = 3.66; 45°: t(11) = 13.83, p < 0.001, d = 3.71].

All groups showed a lower PD_max_ value at the end of the long-term retention and generalization phase than at the beginning [0° group: t(22) = 6.95, p < 0.001, d = 2.64; -90° group: t(22) = 4.73, p < 0.001, d = 1.80; 45° group: t(22) = 7.91, p < 0.001, d = 3.00]. The 0° group did not show a higher FFCF value at the end than at the beginning but the -90° and 45° groups did [0° group: t(11) = -0.25, p = 0.807, d = -0.07; -90° group: t(11) = -4.66, p = 0.001, d = -1.63; 45° group: t(11) = -4.56, p = 0.001, d = -1.41]. To sum up, all participants showed retention or generalization at the beginning. Then, the -90° and 45° groups improved their performances further during the generalization phase, while improvement of the 0° group was only apparent in PD_max_.

We used LMM analysis to assess differences between the groups during the initial phase of retention and generalization (20 trials · 36 participants = 720 observations, ICC = 0.09; Table 1), which revealed statistically significant differences between the groups. The retention performance was better than generalization (p < 0.001 and p = 0.021). Generalization was better for the 45° than the - 90° target (p = 0.017).

**Table 1.**
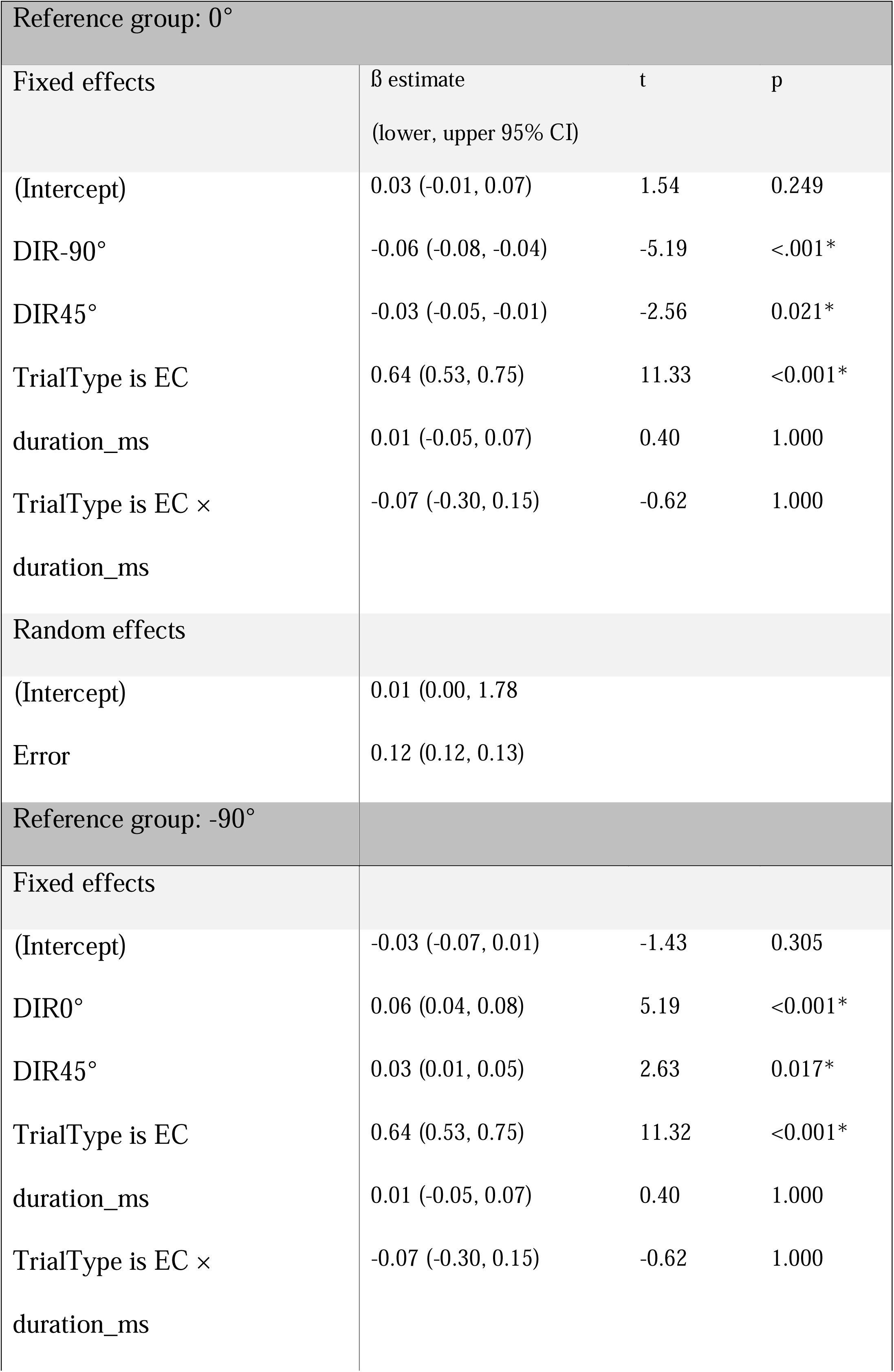
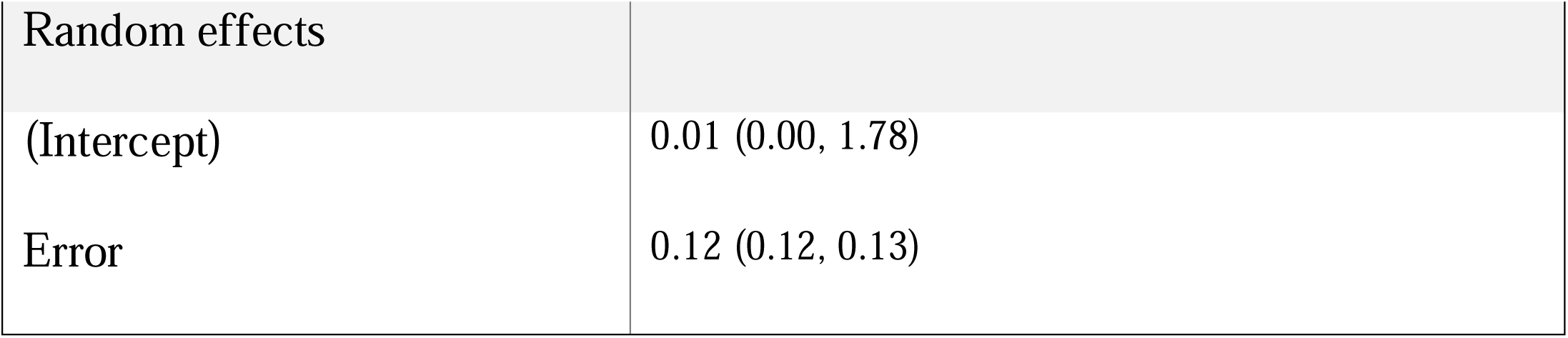
Statistical results of the LMM on the first 20 FF trials of the long-term retention and generalization phases. The asterisks indicate statistical significance after correction for multiple testing.

| Reference group: 0° |  |  |  |
| --- | --- | --- | --- |
| Fixed effects | β estimate<br>(lower, upper 95% CI) | t | p |
| (Intercept) | 0.03 (-0.01, 0.07) | 1.54 | 0.249 |
| DIR-90° | -0.06 (-0.08, -0.04) | -5.19 | <.001* |
| DIR45° | -0.03 (-0.05, -0.01) | -2.56 | 0.021* |
| TrialType is EC | 0.64 (0.53, 0.75) | 11.33 | <0.001* |
| duration_ms | 0.01 (-0.05, 0.07) | 0.40 | 1.000 |
| TrialType is EC ×<br>duration_ms | -0.07 (-0.30, 0.15) | -0.62 | 1.000 |
| Random effects |  |  |  |
| (Intercept) | 0.01 (0.00, 1.78) |  |  |
| Error | 0.12 (0.12, 0.13) |  |  |
| Reference group: -90° |  |  |  |
| Fixed effects |  |  |  |
| (Intercept) | -0.03 (-0.07, 0.01) | -1.43 | 0.305 |
| DIR0° | 0.06 (0.04, 0.08) | 5.19 | <0.001* |
| DIR45° | 0.03 (0.01, 0.05) | 2.63 | 0.017* |
| TrialType is EC | 0.64 (0.53, 0.75) | 11.32 | <0.001* |
| duration_ms | 0.01 (-0.05, 0.07) | 0.40 | 1.000 |
| TrialType is EC ×<br>duration_ms | -0.07 (-0.30, 0.15) | -0.62 | 1.000 |

|  |  |
| --- | --- |
| Random effects |  |
| (Intercept) | 0.01 (0.00, 1.78) |
| Error | 0.12 (0.12, 0.13) |

### 5.4 Two to five muscle synergies are employed during baseline

We used muscle synergy analysis to assess the changes in muscle patterns underlying force field adaptation. First, we extracted muscle synergies from the baseline phase. Over all participants, 3.6 ± 0.6 synergies led to a reconstruction quality (R²) of 0.82 ± 0.04 (Table 2). Figure 4 displays the R² and EMG for every participant, as well as synergies and their reconstruction for one exemplary participant. Across all participants, baseline muscle synergies comprised simultaneous activations of multiple muscles, including mono-articular and bi-articular muscles. Five of the eight baseline muscle synergy clusters were directionally tuned according to a cosine tuning function (Figure 5).

**Figure 4:**
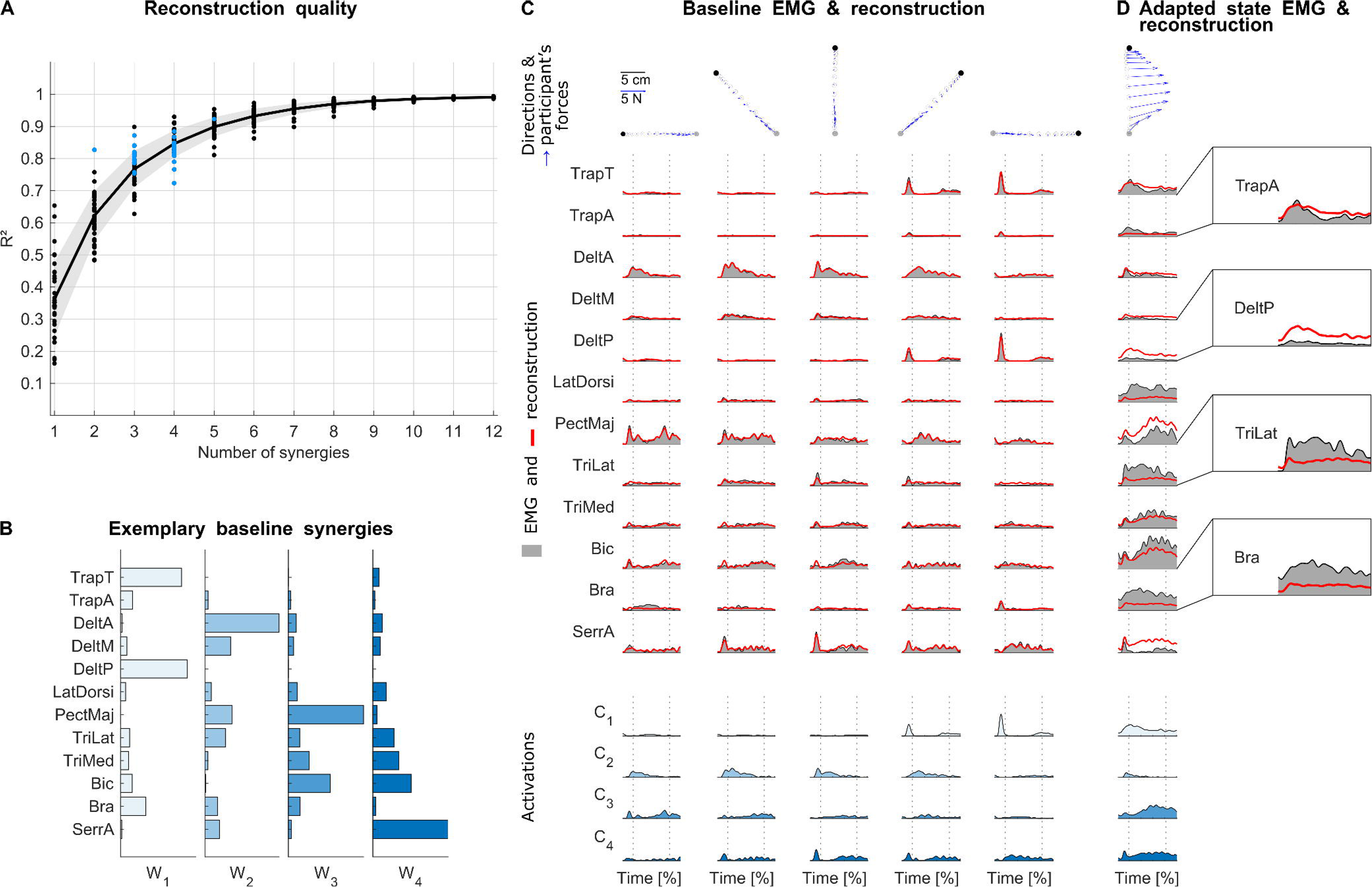
Muscle synergy extraction from baseline. A: Reconstruction quality R² as a function of the number of extracted synergies. The dots represent individual values and the solid, shaded line represents mean and standard deviation. Blue colored dots show the participant-specific selected numbers based on the R²-knee criterion. B: Baseline muscle synergies extracted from one exemplary participant. C Top: Mean trajectory and forces averaged over all trials for every direction during baseline from the same exemplary participant. The distances of the dots illustrating the trajectories display the reaching speed (inter-dot distance corresponds to 8% of the trial duration). C Middle: EMG averaged over all trials for each direction during baseline from the same exemplary participant (gray) and the respective reconstructions (red). C Bottom: Activation functions for every direction in columns. Each activation function belongs to one muscle synergy (e.g., C_1_ belongs to W_1_). D: Trajectories, forces, and EMG averaged over the last 20 FF trials during adaptation for the same participant and the respective reconstruction by the baseline synergies (“baseline fit”). The enlargements show the mismatches in detail for four muscles. The dotted lines in C and D show the time points when the participant left the start point and reached the target point.

**Figure 5:**
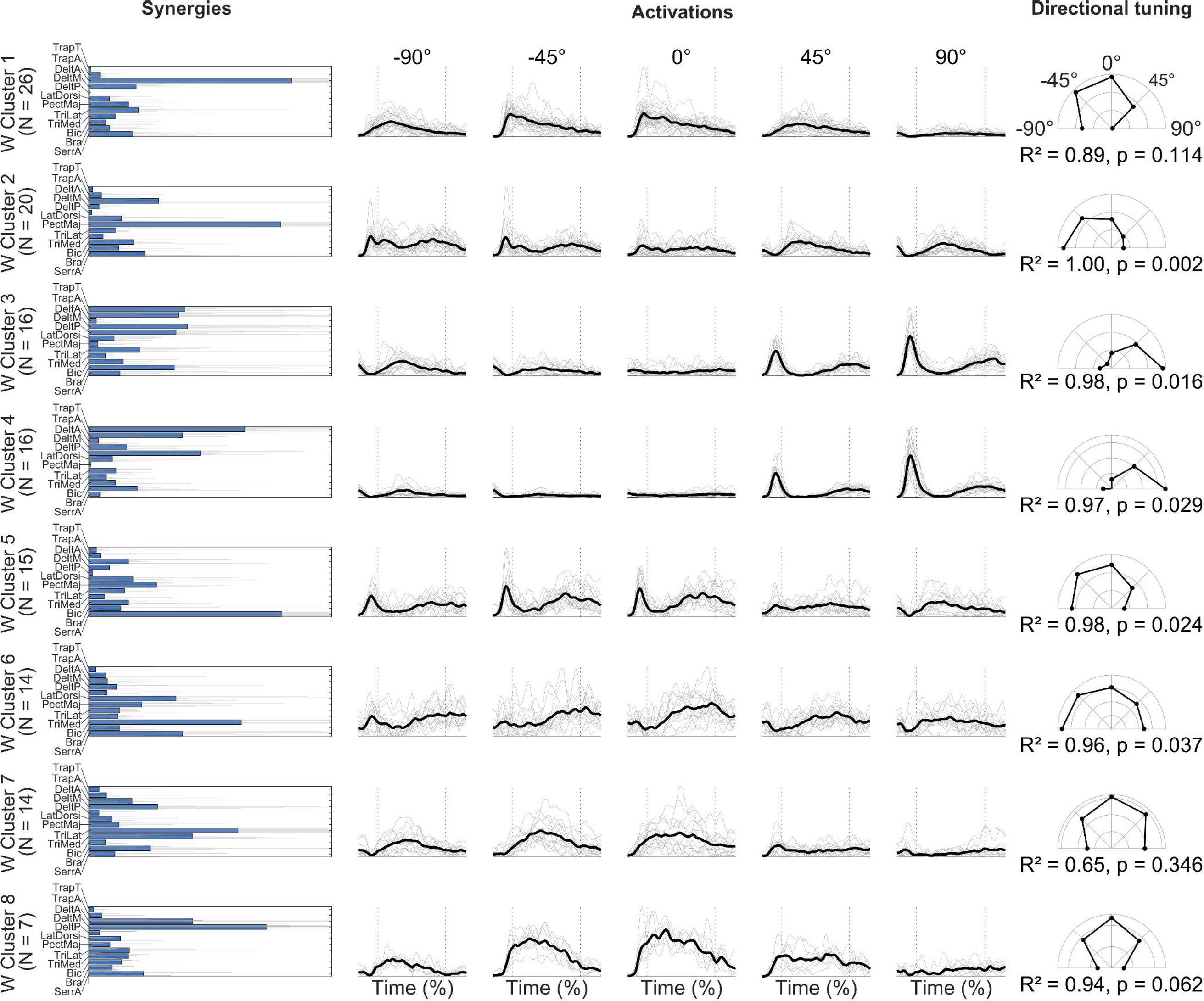
Clustering and cosine fit of the baseline synergies. The left column shows the centroids (filled bars) and the individual synergies in gray solid lines. The middle column shows the mean (black, solid lines) and the individual (gray, solid lines) activation functions separated for the five directions. The dotted lines show the average time points when the participants left the start point and reached the target point. The right column shows the cosine tuning.

**Table 2:**
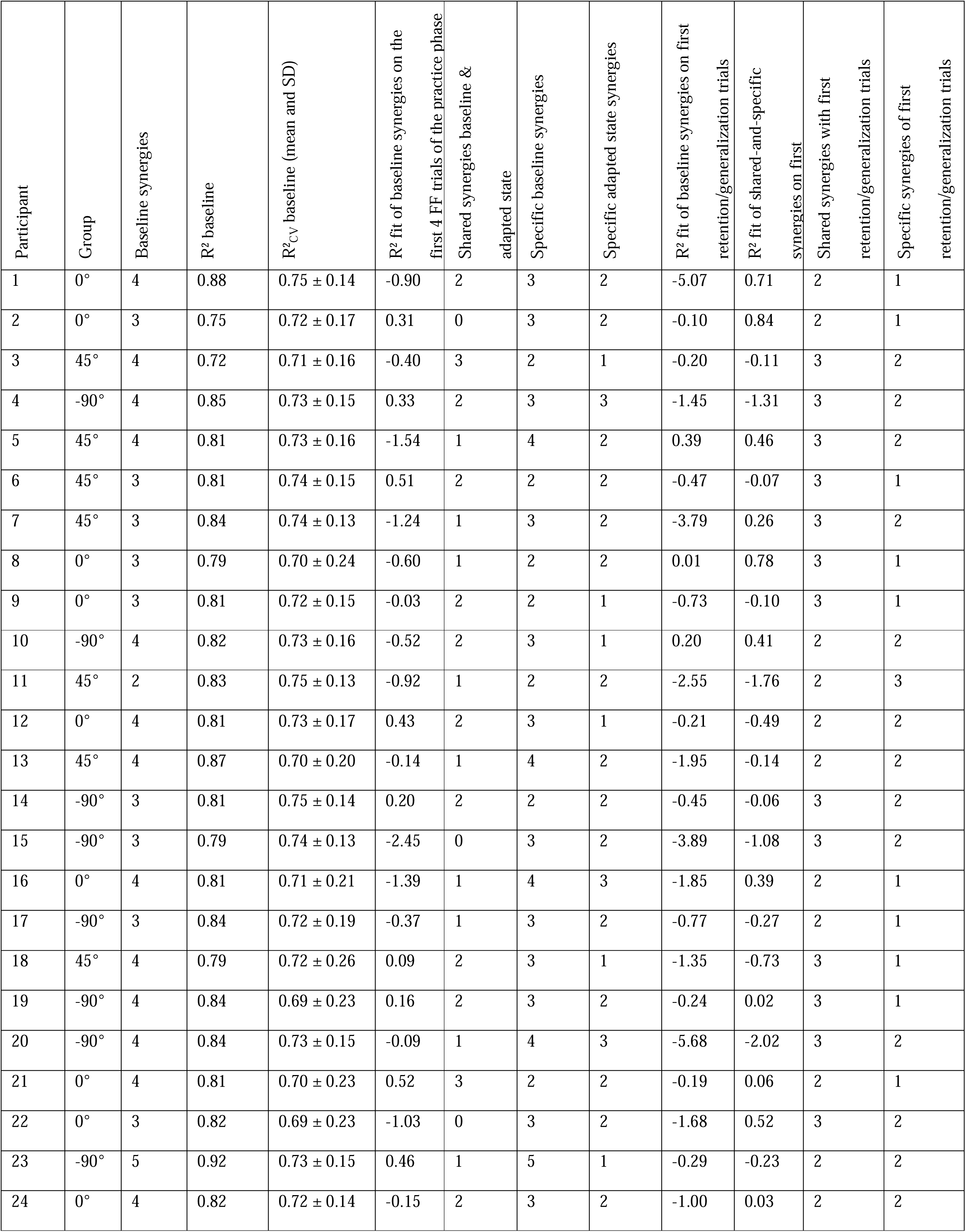
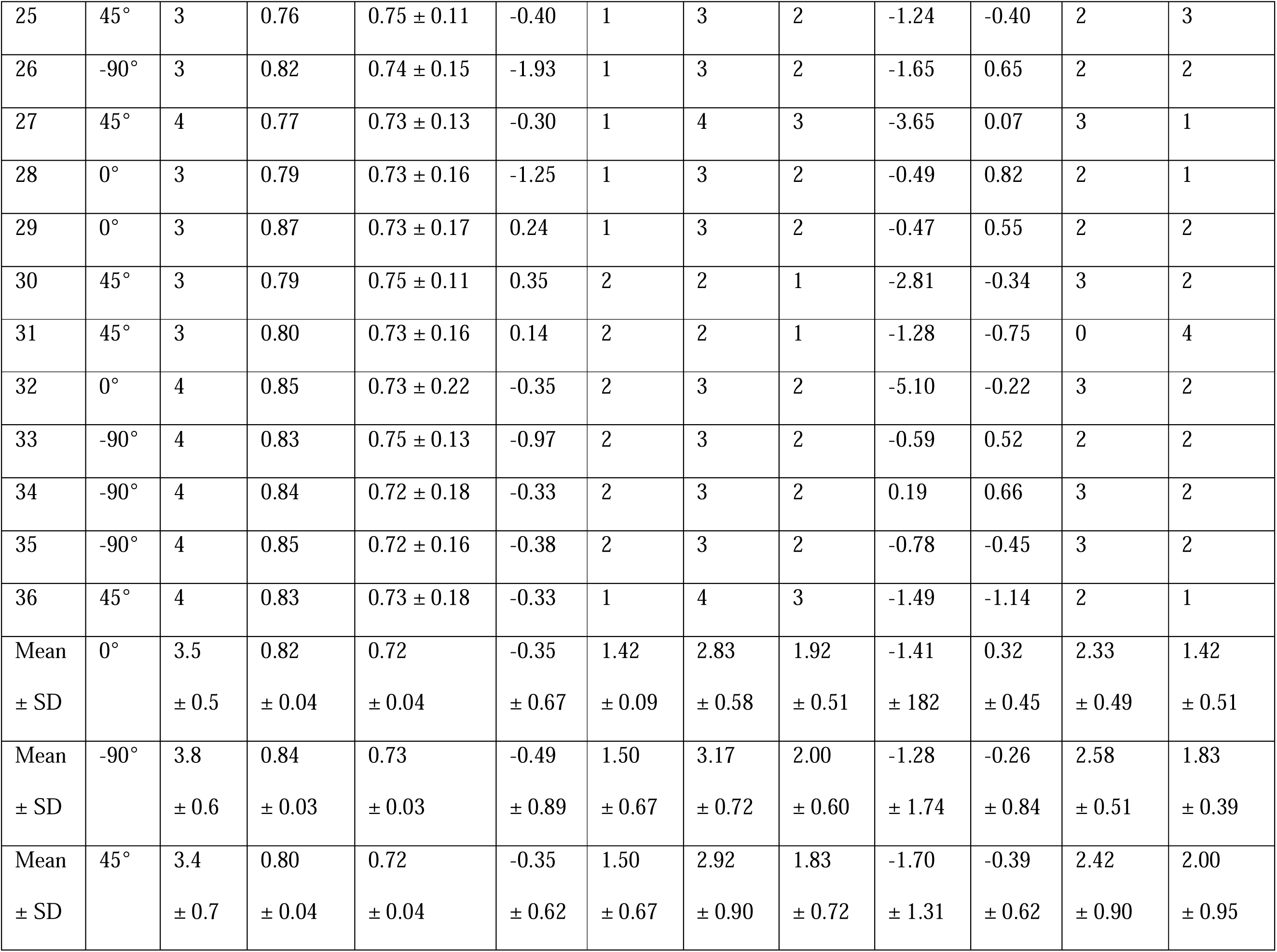
Individual results from the muscle synergy analysis. The last three rows present the results aggregated for the three groups (-90°, 0°, 45°).

### 5.5 The muscle patterns after adaptation cannot be reconstructed by baseline reaching synergies only, and require additional adaptation-specific muscle synergies

We tested if the muscle patterns of force field adaptation can be reconstructed by a combination of baseline reaching synergies (H_Synergies_ 1). The reconstruction quality R² was significantly worse for the adapted state than the cross-validated R²_CV_ of the baseline [t(35) = 7.61, p < 0.001, d = 1.76], with R² values on average 0.98 ± 0.69 lower for every participant (Table 2). This low reconstruction quality, i.e., the mismatch between the reconstructed EMG and the original EMG, is illustrated in the example of Figure 4D. Hence, muscle synergies of unperturbed reaching cannot explain muscle patterns for reaching in a force field, and we reject H_Synergies_ 1.

Subsequently, we tested if specific muscle synergies are required using the bootstrap approach (H_Synergies_ 2). The higher the number of shared synergies, the more subspace dimensions are shared. Across all participants, we found 1.47 ± 0.74 shared synergies but 2.97 ± 0.74 baseline-specific and 1.92 ± 0.60 adapted state-specific synergies. All synergies were clustered across all participants into four shared, seven baseline-specific, and three adapted state-specific synergies clusters (Figures 6-7).

**Figure 6:**
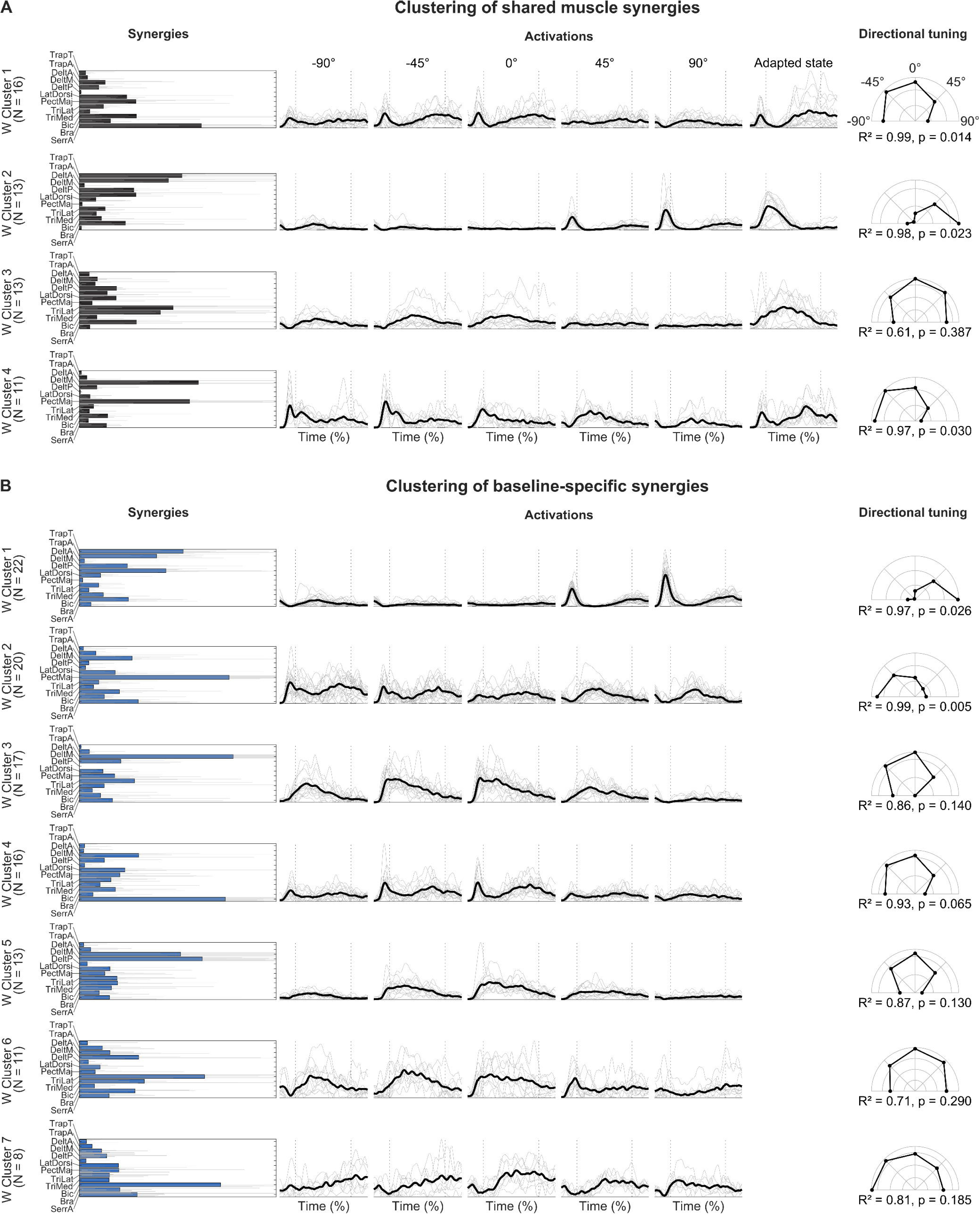
Results of the clustering of the shared-specific muscle synergy extraction of baseline and adapted state. A: Clusters of the shared muscle synergies. B: Clusters of the adapted state-specific muscle synergies. The left column shows the centroids (filled bars) and the individual synergies in gray solid lines. The middle column shows the mean (black, solid lines) and the individual (gray, solid lines) activation functions. The dotted lines show the average time points when the participants left the start point and reached the end point. The right column shows the cosine tuning based on the baseline activation functions.

**Figure 7:**
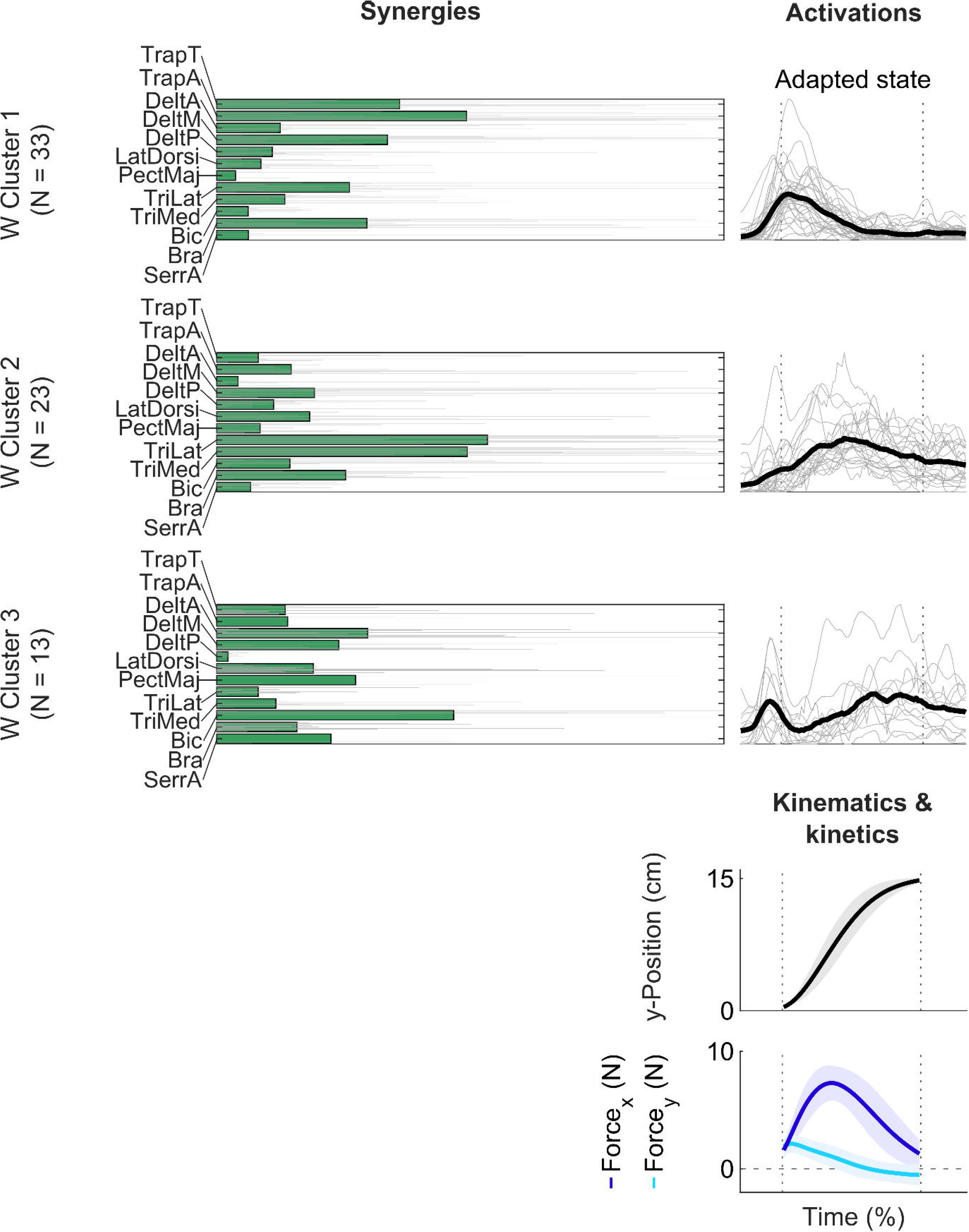
Shared-and-specific synergy extraction and clustering of the adapted state-specific synergies. The left column shows the centroid (filled bars) and the individual synergies in gray solid lines. The right column shows the mean (black, solid lines) and the individual (gray, solid lines) activation functions. Below, the position of the handle in the y-direction is shown as mean (solid black line) and standard deviation (gray shaded) across all participants and the last 20 FF trials. Similarly, the participant’s mean forces in the x- and y-direction are shown as mean (blue solid lines) and standard deviations (blue shaded line). The dotted lines show the average time points when the participants left the start point and reached the end point. The dashed line shows 0N.

Three of the four shared synergies are directionally tuned, two toward the left and one to the right. Two synergies (clusters 1 and 4) show high activation in the middle and end of the adapted state trial and may, therefore, represent a deceleration of the reach. One synergy tuned to the right side is active at the beginning of the adapted state trial, probably to counteract the perturbation. However, the shared synergies do not contribute much to representing adapted state reaching, as an additional 1.92 ± 0.60 adapted state-specific synergies were necessary to describe the muscle pattern in the adapted state.

The baseline-specific synergies and their activation function overall resemble those of the synergies extracted solely from the baseline. The baseline clusters 1, 2, 3, 4, 5, 6, and 7 from the shared-and-specific extraction resemble baseline clusters 3 and 4, 2, 1, 5, 8, 7, and 6 from the baseline extraction only (Figures 5-6). Accordingly, the synergies employed during the baseline are specific to this phase and are not used in the adapted state.

The adapted state-specific synergy cluster 1 (Figure 7) shows high activations of muscles that extend the elbow and the shoulder horizontally, i.e., rotate the arm outward while extending it. This synergy is activated early in the movement, with its peak activity shortly after the start. Cluster 2 shows high activations of TriLat and TriMed, the two main contributors for elbow extension; it is active over the whole trial, with the peak in the middle of the movement. Cluster 3 shows the co-activation of many muscles that act in opposing directions (e.g., DeltA and DeltM, PectMaj and LatDorsi), probably reflecting co-contraction, and especially high activations of Bic, DeltA and DeltM, and PectMaj. These muscles can be seen as antagonists of the aforementioned muscles during the reaching movement toward the 0° target. This synergy shows two peak activations, the first before the beginning of the movement and the second before reaching the target. The three synergies are activated in four phases. After an initial co-contraction (cluster 3), clusters 1 and 2 are activated, leading to forces toward the target (Force_y_) and against the counter-clockwise force field (Force_x_). Interestingly, the two synergies’ activations overlap but have their peaks one after the other; yet Force_x_ has a Gaussian shape without any jitter. Afterward, the arm is decelerated by cluster 3, which is reflected in the negative values of Force_y_.

Figure 8 shows the detailed reconstruction of the adapted state for the exemplary participant with the shared-and-specific synergy extraction. One synergy is shared between the baseline and the adapted state (Figure 8B, W_1_), as obtained by evaluating the principal angle distributions (Figure 8C). There are four baseline-specific synergies (W_2_-W_5_), resembling those of the baseline-only extraction (Figure 4), except that the high PectMaj activity is now present in the shared synergy W_1_. Most interestingly, the adapted state-specific synergies W_6_ and W_7_ show the subsequent but overlapping activation of two synergies which reflect arm extension and outward rotation, probably leading to forces necessary for adapted reaching. This stays in contrast with unperturbed reaching. While one synergy (W_5_) showed agonistic muscle activations for unperturbed 0° reaching, two muscle synergies showed it in adapted state reaching (W_6_ and W_7_). One specialty of this participant is the high activation of the Bic throughout the movement.

**Figure 8:**
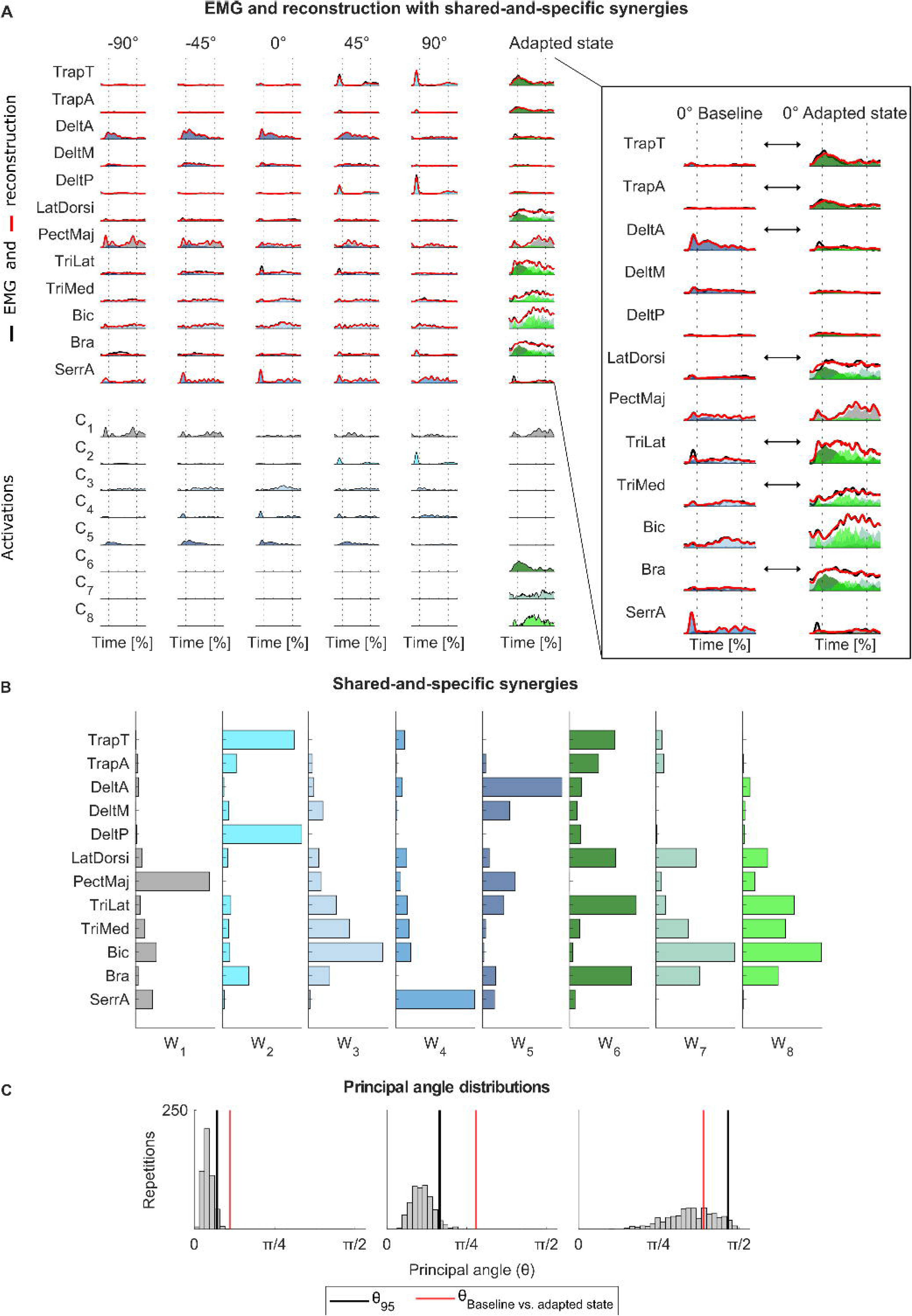
Reconstruction of the baseline and adapted state EMG with the shared-and-specific muscle synergies. A: EMG (solid black lines) and reconstruction (solid red lines) of shared-and-specific synergy extraction of one exemplary participant. The reconstruction is calculated as the sum of the products of the synergies with their respective activation functions. Here, the reconstruction of each product is plotted transparently to illustrate their contributions to the overall reconstruction. The activation functions of the synergies are plotted below. The enlargement shows the reconstruction of the 0° baseline and adapted state in greater detail; arrows guide the reader to substantial differences in synergy activations. B: Muscle synergies. The shared synergy is gray, the baseline-specific synergies in blue shades, and the adapted state-specific synergies in green shades. C: Combined principal angle distributions obtained with the bootstrapping procedure for the exemplary participant. The vertical black line signifies each distribution’s 95th percentile (θ95), and the red line is the principal angle between the baseline and adapted state synergies.

In summary, the adapted state EMG patterns are represented predominantly by adapted state-specific muscle synergies with a four-phasic activation pattern.

### **5.6** Muscle synergies acquired during adaptation facilitate long-term retention and generalization

Kinematics and kinetics indicated that the 0° group showed retention and that the 45° and 90° groups showed generalization, with slightly better values for the 45° group (section 5.3). Therefore, we tested whether muscle synergies can reflect these findings using the shared-and-specific synergies extracted from the baseline and adapted state to reconstruct the muscle patterns of the long-term generalization phase (H_Synergies_ 3).

We found that the shared-and-specific synergies explained the muscle patterns at the beginning of the long-term retention and generalization phase better than the baseline synergies, as all groups showed a significant improvement in the reconstruction quality R² (-90°: W = 78, p < 0.001, d = 1.02; 0°: W = 78, p < 0.001, d = 0.93; 45°: W = 78, p < 0.001, d = 1.16; Table 2, Figure 9A). This means that the muscle patterns during the first trials of the long-term retention and generalization phase can be better explained by the synergies that capture the changes occurring during adaptation (shared-and-specific synergies) than those that do not (baseline synergies). However, there was no difference between the groups regarding the improvement in reconstruction achieved with shared-and-specific synergies with respect to baseline synergies [H(2, 35) = 1.76, p = 0.415, η_p_^2^ = 0.05].

**Figure 9:**
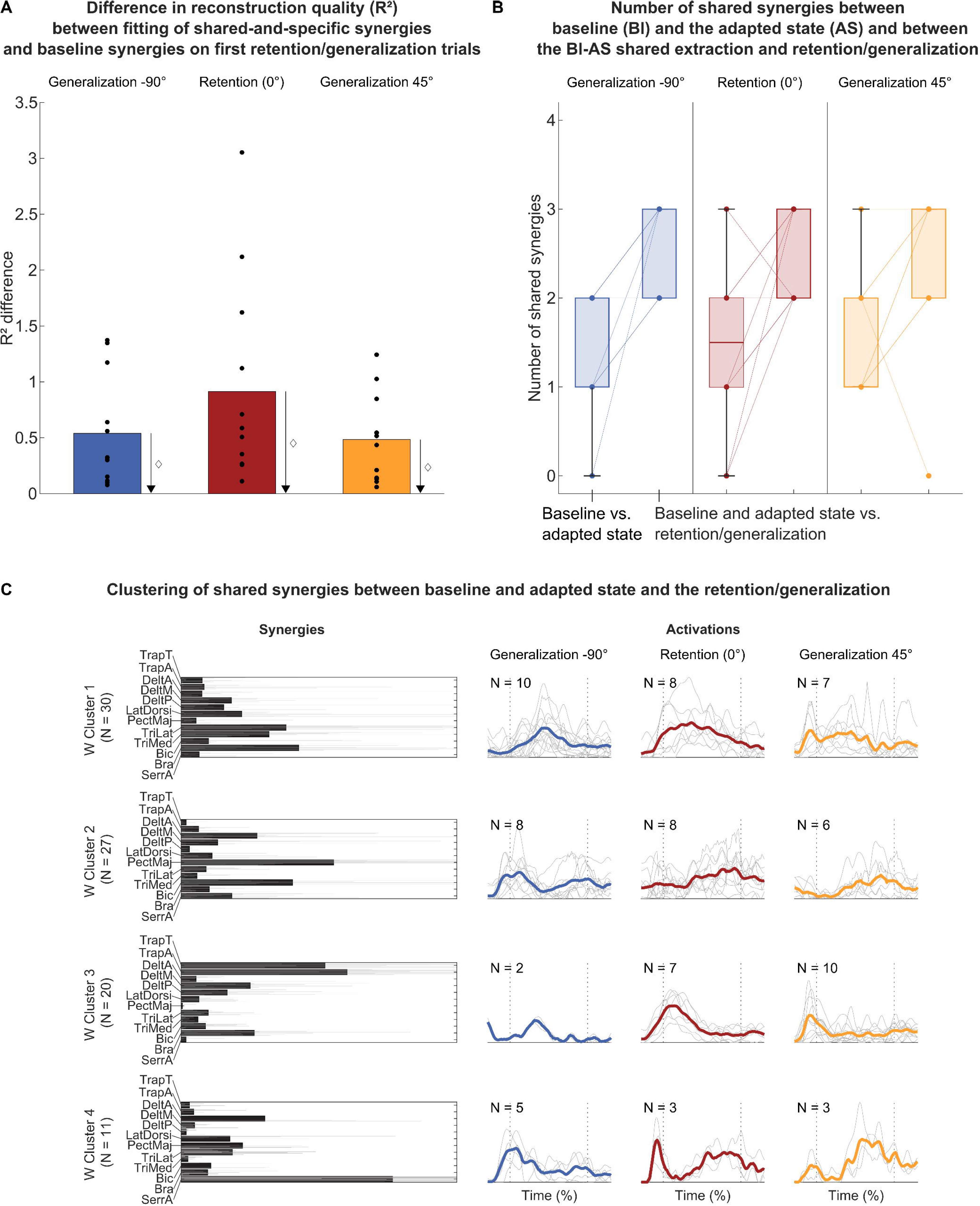
A: Difference in reconstruction quality (R²) between fitting of shared-and-specific and baseline synergies on first retention/generalization trials. Bars represent the mean and dots individual values. The values are presented according to their group. A value larger than zero indicates that shared-specific synergies better reconstructed the EMG of the first retention/generalization trials than baseline synergies. The l1 sign indicates a significant difference from zero B: Comparison of how many synergies are shared between (1) the baseline and the adapted state and (2) the shared-and-specific synergies of the baseline and adapted state together and the first retention/generalization trial. The box chart shows the median and the lower and upper quartiles, and lines show individual values. C: Clustering results of the shared synergies from the shared-and-specific synergy extraction of baseline and adapted state together and the retention/generalization trials. The left column shows the centroid (filled bars) and the individual synergies in gray solid lines. The right columns show the mean (solid lines) and the individual (gray, solid lines) activation functions, separated for the three groups. The dotted lines show the average time points when the participants left the start point and reached the target point.

Next, we investigated the dimensionality and the structure of the muscle synergies that facilitate retention and generalization. This was done by comparing the shared-and-specific synergies extracted from the baseline and adapted state together and the synergies extracted from retention/generalization trials. We found 2.50 ± 0.50 shared dimensions between the subspaces spanned by the baseline and adapted state synergies and retention/generalization synergies. Hence, more synergies are shared in this case than between baseline and adapted state synergies (Figure 9B). Accordingly, the higher number of shared dimensions indicates that the muscle synergies reflect the findings of retention and generalization in the kinematic and kinetic variables. The acquired structural changes in the synergies that occurred during adaptation may facilitate retention and generalization. Figure 9C shows the clusters of the shared synergies between the baseline and adapted state synergies and the retention/generalization synergies. The first three resemble the adapted state-specific synergies reported in section 5.5 (clusters 1, 2, and 3 in Figure 9C resemble clusters 2, 3, and 1 in Figure 7). Furthermore, for the retention, we observe the described four-phasic pattern again. Also, the 45° group shows a similar four-phasic pattern, with the same synergies activated one after another, as the 0° retention group. The -90° group differs from the 0° and 45° groups. Cluster 1 is activated later in the movement and cluster 2 is activated earlier, probably accelerating the arm toward the target instead of decelerating, and only two participants show synergies for cluster 3. Still, we observe that two muscle synergies are activated in an overlapping matter with subsequent activity peaks when the handle moves between the start and stop targets, just like the 0° and 45° groups.

However, the differences between the groups observed with PD_max_ and FFCF are not well reflected in the modular structure, as we found no differences regarding the reconstruction quality of the baseline and adapted state shared-and-specific synergies on the first retention/generalization trials or the number of specific synergies between the -90° and the 45° groups. Hence, we reject H_Synergies_ 3.

## 6 Discussion

This study investigated the relationship between muscle synergies and force field adaptation, retention, and generalization. Our findings show that adaptation involves structural changes in muscle synergies compared to reaching in unperturbed conditions, alongside a novel four-phasic synergy activation pattern. Moreover, these structural changes and activation patterns likely facilitate retention and generalization, as the same synergies and their activation patterns are also reflected there.

### 6.1 Participants adapted to the force field, showed retention to the 0° target, and generalization to the 45° and -90° targets

Participants adapted to the force field during their first exposure, washed out after its removal, and re-adapted and generalized when re-exposed (H_Task_ 1). Also, we found that generalization decreased with distance from the practiced movement direction (H_Task_ 2). These results align with related force field studies (Shadmehr and Mussa-Ivaldi, 1994; Brashers-Krug et al., 1996; Gandolfo et al., 1996; Rezazadeh and Berniker, 2019; Mathew et al., 2021). The fast re-adaptation for the 0°, i.e., the same direction as during the first exposure, has been previously described as the “savings” effect (Brashers-Krug et al., 1996; Shadmehr and Brashers-Krug, 1997; Mathew et al., 2021). However, based on the aforementioned studies, we expected no generalization to the -90° target at the beginning of the long-term retention and generalization phase. This disagreement may stem from participants already being exposed to the -90° FF condition during the short-term retention and generalization test, as adaptation starts from the very first trial (Joiner et al., 2017). The adaptation, washout, savings, and generalization we identified allowed us to examine the underlying modular structure of the muscle patterns.

### 6.2 Reaching in an environment with altered dynamics requires structural changes to the muscle synergies for unperturbed reaching

We first extracted muscle synergies from the planar, center-out reaching movements in the null field during the baseline. The number, composition, and tuning of the extracted baseline synergies are in accordance with the literature (d’Avella et al., 2006; Muceli et al., 2010).

Since a possible mechanism for reaching in a force field could be to combine the synergies employed in the baseline, we hypothesized that the muscle patterns of force field adaptation can be reconstructed by a combination of baseline reaching synergies (H_Synergies_ 1). However, our results indicate this is not the case, and force field adaptation requires structural changes in muscle synergies (H_Synergies_ 2). We found adapted state-specific synergies activated in a four-phasic pattern. First, a synergy reflecting co-contraction is active. Then, there is an early onset of a synergy reflecting arm extension and outward rotation, which is active until halfway through the trial until a second synergy overtakes, mainly reflecting triceps activity and, thus, elbow extension. At the trial end, the movement is decelerated and stabilized through a synergy of antagonistic muscles.

Reaching movements have been found to follow a triphasic pattern of muscle activation, leading to acceleration, deceleration, and damping of the movement (Wadman et al., 1979; Happee, 1992; Flanders et al., 1994). This pattern is generally also reflected in the activation of muscle synergies, showing an interplay between agonistic and antagonistic muscle synergies (d’Avella et al., 2006; Chiovetto et al., 2013). These observations regarding EMG and muscle synergies hold for baseline reaching in our study. Also, when adapted, participants employed a triphasic pattern of muscle activity as previously described (Thoroughman and Shadmehr, 1999; Darainy and Ostry, 2008). However, the novelty of our findings is that the muscle activity used to move the hand to the target is realized through two synergies with overlapping activations, peaking sequentially. Interestingly, the transition between the two synergies is seamless, as the force curves are bell-shaped and smooth. We provide a novel characterization of changes in the synergistic organization of many muscles after adaptation to a perturbing force field. Nevertheless, our findings align with literature examining activities in a few muscles, when looking at the muscle activity in our study without looking at the muscle synergies. We also observed early activity in these muscles, probably reflecting co-contraction (Milner and Franklin, 2005; Darainy and Ostry, 2008), and the muscles for reaching forward and counteracting the force field are already active early in the movement (Thoroughman and Shadmehr, 1999; Albert and Shadmehr, 2016).

Huang et al. (2012) showed that muscle activity is reduced to minimal levels even after a plateau in kinematic- and kinetic-dependent variables. Accordingly, the CNS might optimize effort while adapting, which leads to specific muscle synergies. The synergy that reflects arm extension with high triceps activation is sparse, and may reflect this tendency to realize movements with less effort. However, there is a lack of clarity over the process of how the structural changes happen, i.e., if adaptation expands the subspaces spanned by baseline synergies by learning new synergies or if adapted state-specific subspaces are the results of other control processes not requiring new synergies. This unmet need motivates future studies.

Force field adaptation with a muscle synergy perspective has received little research attention. Oscari et al. (2016) found that moving a joystick in a force field involves two additional muscle synergies during adaptation. This supports the notion of structural changes during force field adaptation, even though their results may be limited. Sampling a single direction may limit the validity of extracted synergies (Steele et al., 2015), characterizing only acceleration and deacceleration patterns (Chiovetto et al., 2013) and neglecting the versatile use of baseline reaching synergies to different directions. In contrast to our findings, a structural change in muscle synergies has not previously been reported during visuomotor rotations (Gentner et al., 2013; De Marchis et al., 2018; Severini and Zych, 2020). Here, muscle synergies extracted from the baseline could reconstruct the muscle patterns during adaptation and generalization, indicating structural robustness. Although force field and visuomotor adaptation are related motor adaptation paradigms, there are some differences, especially when the latter is done under isometric conditions. In contrast to isometric reaching, in dynamic reaching the joint angles and muscle length change throughout the movement; and the force exerted by the participant on the handle depends on the joint angle, muscle length, and their derivatives (Bizzi et al., 1991; Giszter et al., 1993; Mussa-Ivaldi et al., 1994; Shadmehr and Wise, 2005). It is, therefore, plausible that force field adaptation requires muscle synergies to be appropriate and effort-optimized at all joint angle configurations used during the reach, and that a re- aiming strategy by tuning baseline synergies does not suffice.

### 6.3 Retention and generalization are represented in the modular structure after adaptation

We found that retention and generalization muscle patterns can be better explained with synergies extracted from both baseline and adaptation than from baseline synergies alone. Therefore, the structural changes acquired during adaptation are re-used during retention and generalization. The retention (0° target) muscle pattern is well described by the shared synergies. Hence, our results show that muscle synergies and their activation timing are re-used when re-exposed to the same force field, presumably allowing a fast re-adaptation. This suggests that the shared muscle synergies represent a mechanism capturing the savings effect at a modular level (Brashers-Krug et al., 1996; Shadmehr and Brashers-Krug, 1997; Mathew et al., 2021).

However, contrary to our hypothesis H_Synergies_ 3, the different amount of generalization between the - 90° and 45° directions is not evident in the reconstruction quality, the number of shared synergies, or the number of direction-specific synergies. Accordingly, the structural changes in muscle synergies through adaptation reflect retention and generalization but not the differences between the 0°, -90°, and 45° groups. The observed structural changes through adaptation probably represent a general, i.e., direction-invariant, coordinative solution to the force field perturbation. This supports the notion that muscle synergies represent low-dimensional, modular control and are re-used in reaching to multiple directions (d’Avella et al., 2006), yet motivates future studies to further investigate the relationship between muscle coordination and task-level generalization performances. For example, future studies may investigate the trial-by-trial changes separately for the -90° and 45° directions to investigate possible mechanisms related to different generalization speeds.

### 6.4 Limitations

Baseline reaching directions comprised a semi-circle, and only center-out movements were analyzed. This restricts the possible subspaces and, thus, potentially shared subspace dimensions with the adapted state. Also, all participants adapted to the 0° target first, so future studies may generalize our findings with different directions. Due to the limited number of movements, we cannot conclusively state whether the CNS acquired new synergies during adaptation, or if these synergies are stored in the CNS but were not recruited in the baseline. Furthermore, future studies may investigate how the structural changes in muscle synergies evolve with practice.

### 6.5 Conclusions

This study found that reaching in an environment with altered dynamics requires structural changes and novel activation patterns in muscle synergies. These structural changes facilitate retention when re-adapting to the same direction, and generalization when adapting to new directions. Thus, our results provide new insights into how force field adaptation, retention, and spatial generalization are represented at the level of muscular coordination.

## 7 Author contributions

- MH and TS designed the research.
- MH performed the research.
- MH, DJB, MR, ADA, and TS analyzed the data.
- MH, DJB, MR, ADA, and TS wrote the paper.

## 8 Funding

### N/A

### 9 Data availability statement

The raw data supporting the conclusions of this article will be made available by the authors without undue reservation.

## References

1. Ahmad I, Ansari F, Dey UK (2013) Power Line Noise Reduction In Ecg By Butterworth Notch Filters: A Comparative Study. IJECIERD 3:65–74.

2. Albert ST, Shadmehr R (2016) The Neural Feedback Response to Error As a Teaching Signal for the Motor Learning System. J Neurosci 36:4832–4845.

3. Allen JL, Kesar TM, Ting LH (2019) Motor module generalization across balance and walking is impaired after stroke. J Neurophysiol 122:277–289.

4. Anwar MN, Tomi N, Ito K (2011) Motor imagery facilitates force field learning. Brain Res 1395:21– 29.

5. Bach MM, Daffertshofer A, Dominici N (2021) Muscle Synergies in Children Walking and Running on a Treadmill. Front Hum Neurosci 15.

6. Bernstein NA (1967) The co-ordination and regulation of movements. Oxford: Pergamon Press.

7. Bizzi E, Cheung VCK, d’Avella A, Saltiel P, Tresch M (2008) Combining modules for movement. Brain Res Rev 57:125–133.

8. Bizzi E, Mussa-Ivaldi FA, Giszter S (1991) Computations underlying the execution of movement: a biological perspective. Science 253:287–291.

9. Brambilla C, Russo M, d’Avella A, Scano A (2023) Phasic and tonic muscle synergies are different in number, structure and sparseness. Hum Mov Sci 92:103148.

10. Brashers-Krug T, Shadmehr R, Bizzi E (1996) Consolidation in human motor memory. Nature 382:252–255.

11. Carey HD, Liss DJ, Allen JL (2021) Young adults recruit similar motor modules across walking, turning, and chair transfers. Physiol Rep 9:e15050.

12. Cheung VCK, d’Avella A, Bizzi E (2009) Adjustments of Motor Pattern for Load Compensation Via Modulated Activations of Muscle Synergies During Natural Behaviors. J Neurophysiol 101:1235–1257.

13. Cheung VCK, d’Avella A, Tresch MC, Bizzi E (2005) Central and Sensory Contributions to the Activation and Organization of Muscle Synergies during Natural Motor Behaviors. J Neurosci 25:6419–6434.

14. Chiovetto E, Berret B, Delis I, Panzeri S, Pozzo T (2013) Investigating reduction of dimensionality during single-joint elbow movements: a case study on muscle synergies. Front Comput Neurosci 7.

15. Cohen J (1988) Statistical power for the behavioural sciences, 2nd ed. Hilsdale: Lawrence Erlbaum Associates.

16. d’Avella A (2016) Modularity for Motor Control and Motor Learning. In: Progress in Motor Control: Theories and Translations (Laczko J, Latash ML, eds), pp 3–19 Advances in Experimental Medicine and Biology. Cham: Springer International Publishing.

17. d’Avella A, Bizzi E (2005) Shared and specific muscle synergies in natural motor behaviors. PNAS 102:3076–3081.

18. d’Avella A, Portone A, Fernandes L, Lacquaniti F (2006) Control of fast-reaching movements by muscle synergy combinations. J Neurosci 26:7791–7810.

19. d’Avella A, Saltiel P, Bizzi E (2003) Combinations of muscle synergies in the construction of a natural motor behavior. Nat Neurosci 6:300–308.

20. Darainy M, Ostry DJ (2008) Muscle cocontraction following dynamics learning. Exp Brain Res 190:153–163.

21. De Marchis C, Di Somma J, Zych M, Conforto S, Severini G (2018) Consistent visuomotor adaptations and generalizations can be achieved through different rotations of robust motor modules. Sci Rep 8:12657.

22. Diedrichsen J, White O, Newman D, Lally N (2010) Use-Dependent and Error-Based Learning of Motor Behaviors. J Neurosci 30:5159–5166.

23. Flanders M, Pellegrini JJ, Soechting JF (1994) Spatial/temporal characteristics of a motor pattern for reaching. J Neurophysiol 71:811–813.

24. Franklin DW, Osu R, Burdet E, Kawato M, Milner TE (2003) Adaptation to Stable and Unstable Dynamics Achieved By Combined Impedance Control and Inverse Dynamics Model. J Neurophysiol 90:3270–3282.

25. Franklin S, Franklin DW (2021) Feedback Gains modulate with Motor Memory Uncertainty. Neurons behav data anal theory 5:1–28.

26. Gandolfo F, Mussa-Ivaldi FA, Bizzi E (1996) Motor learning by field approximation. PNAS 93:3843–3846.

27. Gentner R, Edmunds T, Pai DK, d’Avella A (2013) Robustness of muscle synergies during visuomotor adaptation. Front Comput Neurosci 7.

28. Ghez C, Krakauer JW, Sainburg R, Ghilardi M (1999) Spatial representations and internal models of limb dynamics in motor learning. In: The Cognitive Neurosciences, 2nd ed., pp 501–514. Cambridge: The MIT Press.

29. Giszter SF (2015) Motor primitives—new data and future questions. Curr Opin Neurobiol 33:156– 165.

30. Giszter SF, Mussa-Ivaldi FA, Bizzi E (1993) Convergent force fields organized in the frog’s spinal cord. J Neurosci 13:467–491.

31. Golub GH, Van Loan CF (1989) Matrix Computations, 2nd ed. Baltimore: Johns Hopkins University Press.

32. Happee R (1992) Goal-directed arm movements: I. Analysis of EMG records in shoulder and elbow muscles. J Electromyogr Kinesiol 2:165–178.

33. Heald JB, Franklin DW, Wolpert DM (2018) Increasing muscle co-contraction speeds up internal model acquisition during dynamic motor learning. Sci Rep 8:1–11.

34. Hermens HJ, Freriks B, Disselhorst-Klug C, Rau G (2000) Development of recommendations for SEMG sensors and sensor placement procedures. J Electromyogr Kinesiol 10:361–374.

35. Herzog M, Focke A, Maurus P, Thürer B, Stein T (2022) Random Practice Enhances Retention and Spatial Transfer in Force Field Adaptation. Front Hum Neurosci 16.

36. Holm S (1979) A simple sequentially rejective multiple test procedure. Scand J Stat 2:65–70.

37. Hox J, Moerbeek M, van de Schoot,Rens (2017) Multilevel AnalysisL: Techniques and Applications, Third Edition, 3rd ed. London: Routledge.

38. Huang HJ, Kram R, Ahmed AA (2012) Reduction of Metabolic Cost during Motor Learning of Arm Reaching Dynamics. J Neurosci 32:2182–2190.

39. Ippersiel P, Preuss R, Fillion A, Jean-Louis J, Woodrow R, Zhang Q, Robbins SM (2021) Inter-joint coordination and the flexion-relaxation phenomenon among adults with low back pain during bending. Gait Posture 85:164–170.

40. Joiner WM, Sing GC, Smith MA (2017) Temporal specificity of the initial adaptive response in motor adaptation. PLOS Comput Biol 13:e1005438.

41. Joiner WM, Smith MA (2008) Long-Term Retention Explained by a Model of Short-Term Learning in the Adaptive Control of Reaching. J Neurophysiol 100:2948–2955.

42. Krakauer JW, Mazzoni P (2011) Human sensorimotor learning: adaptation, skill, and beyond. Curr Opin Neurobiol 21:636–644.

43. Lee DD, Seung HS (1999) Learning the parts of objects by non-negative matrix factorization. Nature 401:788–791.

44. Lee DD, Seung HS (2001) Algorithms for Non-negative Matrix Factorization. In: Advances in Neural Information Processing Systems 13 (Leen TK, Dietterich TG, Tresp V, eds), pp 556–562. MIT Press.

45. Mathew J, Lefèvre P, Crevecoeur F (2021) Savings in Human Force Field Learning Supported by Feedback Adaptation. eNeuro 8:ENEURO.0088-21.2021.

46. Milner T, Franklin DW (2005) Impedance control and internal model use during the initial stage of adaptation to novel dynamics in humans. J Physiol 567:651–664.

47. Muceli S, Boye AT, d’Avella A, Farina D (2010) Identifying Representative Synergy Matrices for Describing Muscular Activation Patterns During Multidirectional Reaching in the Horizontal Plane. J Neurophysiol 103:1532–1542.

48. Mussa-Ivaldi FA (1999) Modular features of motor control and learning. Curr Opin Neurobiol 9:713–717.

49. Mussa–Ivaldi FA, Bizzi E (2000) Motor learning through the combination of primitives. Phil Trans R Soc Lond B 355:1755–1769.

50. Mussa-Ivaldi FA, Giszter SF, Bizzi E (1994) Linear combinations of primitives in vertebrate motor control. PNAS 91:7534–7538.

51. Oldfield RC (1971) The assessment and analysis of handedness: The Edinburgh inventory. Neuropsychologia 9:97–113.

52. Oscari F, Finetto C, Kautz SA, Rosati G (2016) Changes in muscle coordination patterns induced by exposure to a viscous force field. J Neuroeng Rehabil 13:58.

53. Peri E, Xu L, Ciccarelli C, Vandenbussche NL, Xu H, Long X, Overeem S, van Dijk JP, Mischi M (2021) Singular Value Decomposition for Removal of Cardiac Interference from Trunk Electromyogram. Sensors 21:573.

54. Perotto AO (2011) ANATOMICAL GUIDE FOR THE ELECTROMYOGRAPHER: The Limbs and Trunk, 5th ed. Springfield: Charles C Thomas Publisher.

55. Rezazadeh A, Berniker M (2019) Force field generalization and the internal representation of motor learning. PLOS ONE 14:e0225002.

56. Russell DM, Haworth JL (2014) Walking at the preferred stride frequency maximizes local dynamic stability of knee motion. J Biomech 47:102–108.

57. Russo M, Scano A, Brambilla C, d’Avella A (2024) SynergyAnalyzer: A Matlab toolbox implementing mixed-matrix factorization to identify kinematic-muscular synergies. Comput Methods Programs Biomed 251:108217.

58. Scheidt RA, Reinkensmeyer DJ, Conditt MA, Rymer WZ, Mussa-Ivaldi FA (2000) Persistence of motor adaptation during constrained, multi-joint, arm movements. J Neurophysiol 84:853– 862.

59. Severini G, Zych M (2020) Characterization of the Adaptation to Visuomotor Rotations in the Muscle Synergies Space. Front Bioeng Biotechnol 8:605.

60. Shadmehr R (2004) Generalization as a behavioral window to the neural mechanisms of learning internal models. Hum Mov Sci 23:543–568.

61. Shadmehr R (2017) Learning to Predict and Control the Physics of Our Movements. J Neurosci 37:1663–1671.

62. Shadmehr R, Brashers-Krug T (1997) Functional Stages in the Formation of Human Long-Term Motor Memory. J Neurosci 17:409–419.

63. Shadmehr R, Mussa-Ivaldi FA (1994) Adaptive representation of dynamics during learning of a motor task. J Neurosci 14:3208–3224.

64. Shadmehr R, Wise SP (2005) The Computational Neurobiology of Reaching and Pointing: A Foundation for Motor Learning. Cambridge: MIT Press.

65. Sing GC, Joiner WM, Nanayakkara T, Brayanov JB, Smith MA (2009) Primitives for motor adaptation reflect correlated neural tuning to position and velocity. Neuron 64:575–589.

66. Steele KM, Tresch MC, Perreault EJ (2015) Consequences of biomechanically constrained tasks in the design and interpretation of synergy analyses. J Neurophysiol 113:2102–2113.

67. Stockinger C, Thürer B, Focke A, Stein T (2015) Intermanual transfer characteristics of dynamic learning: direction, coordinate frame, and consolidation of interlimb generalization. J Neurophysiol 114:3166–3176.

68. Stone M (1974) Cross-Validatory Choice and Assessment of Statistical Predictions. Journal of the Royal Statistical Society Series B: Statistical Methodology 36:111–133.

69. Sylos-Labini F, La Scaleia V, Cappellini G, Fabiano A, Picone S, Keshishian ES, Zhvansky DS, Paolillo P, Solopova IA, d’Avella A, Ivanenko Y, Lacquaniti F (2020) Distinct locomotor precursors in newborn babies. PNAS 117:9604–9612.

70. Thoroughman KA, Shadmehr R (1999) Electromyographic Correlates of Learning an Internal Model of Reaching Movements. J Neurosci 19:8573–8588.

71. Thoroughman KA, Shadmehr R (2000) Learning of action through adaptive combination of motor primitives. Nature 407:742–747.

72. Ting LH, Macpherson JM (2005) A limited set of muscle synergies for force control during a postural task. J Neurophysiol 93:609–613.

73. Tresch MC, Saltiel P, Bizzi E (1999) The construction of movement by the spinal cord. Nat Neurosci 2:162–167.

74. Wadman WJ, Denier Van der Gon J, Geuze RH, Mol CR (1979) Control of fast goal-directed arm movements. J Hum Mov Stud 5:3–17.

75. Wagner MJ, Smith MA (2008) Shared Internal Models for Feedforward and Feedback Control. J Neurosci 28:10663–10673.

76. Wolpert DM, Kawato M (1998) Multiple paired forward and inverse models for motor control. Neural Netw 11:1317–1329.

